# Segmental Duplications Drive the Evolution of Accessory Regions in a Major Crop Pathogen

**DOI:** 10.1101/2023.06.07.544053

**Authors:** A.C. van Westerhoven, C. Aguilera-Galvez, G. Nakasato-Tagami, X. Shi-Kunne, E. Martinez de la Parte, E. Chavarro-Carrero, H.J.G. Meijer, A. Feurtey, N. Maryani, N. Ordóñez, H. Schneiders, K. Nijbroek, A. H. J. Wittenberg, R. Hofstede, F. García-Bastidas, E.H. Sørensen, R. Swennen, A. Drenth, E.H. Stukenbrock, G.H.J. Kema, M.F. Seidl

**Author notes:** Corresponding authors: Gert H.J. Kema and Michael F. Seidl **Email:**. Authors contributed equally (G.H.J. Kema, M.F. Seidl). **Author Contributions:** ACW collected data and performed the analyses. EMP, CAG, ECC, RH, AW, HS, FGB, NM, NO, EHS and GNT collected data. EHS, AF, KN and XSK contributed to the analyses. ACW and wrote the manuscript with input from HJGM, GHJK, and MFS. All authors contributed to writing and editing the manuscript. GHJK, RS, and MFS conceived and supervised the project. **Competing Interest Statement:** The authors declare no competing interests.

## Abstract

- Many pathogens evolved compartmentalized genomes with conserved core and variable accessory regions which carry effector genes mediating virulence. The fungal plant pathogen *Fusarium oxysporum* has such accessory regions often spanning entire chromosomes. The presence of specific accessory regions influences the host range, and horizontal transfer of some accessory regions can modify the pathogenicity of the receiving strain. However, understanding how these accessory regions evolve in strains that infect the same host remains limited.
- Here, we define the pan-genome of 69 diverse *Fusarium* strains that cause Fusarium wilt of banana, a significant constraint to global banana production. In this diverse panel of *Fusarium* strains infecting banana, we analyzed the diversity and evolution of the accessory regions.
- Accessory regions in *Fusarium* strains infecting the same banana cultivar are highly diverse, and we could not identify any shared genomic regions and in planta induced effectors. We demonstrate that segmental duplications drive the evolution of accessory regions. Furthermore, we show that recent segmental duplications and aneuploidy occur specifically in accessory chromosomes and cause the expansion of accessory regions in *F. oxysporum*.
- Taken together we conclude that extensive recent duplications drive the evolution of accessory regions in *Fusarium*, which contribute to the evolution of virulence.

## Introduction

The interaction between fungal plant pathogens and their hosts drives rapid evolution (Möller & Stukenbrock, 2017). Plant hosts have evolved sophisticated immune systems to detect pathogens, while adapted pathogens, in turn, secrete effectors to deregulate immune responses and to support host colonization (Rovenich *et al*., 2014; Cook *et al*., 2015). To facilitate this co-evolutionary cycle, many filamentous plant pathogens evolved compartmentalized genomes where effector genes are localized in distinct genomic regions (Dong *et al*., 2015; Torres *et al*., 2020). These effector-rich compartments show extensive genetic variation and are enriched for repetitive sequences such as transposable elements (Raffaele & Kamoun, 2012; Dong *et al*., 2015; Seidl & Thomma, 2017; Möller & Stukenbrock, 2017; Torres *et al*., 2020). This spatial separation is often referred to as the ‘two-speed’ genome (Dong *et al*., 2015) and is thought to facilitates the rapid diversification of effector gene repertoires to enable pathogens to evade host resistances and to enable host jumps (Sánchez-Vallet *et al*., 2018).

*Fusarium oxysporum* is a genetically diverse fungal species complex that consists of three clades that can be further separated into different lineages, recently reclassified as separate species (Maryani et al., 2019). Members of this species complex can collectively infect more than 100 economically important crops. In contrast, individual strains typically only infect a single host plant (Edel-Hermann & Lecomte, 2019). *Fusarium oxysporum* evolved a two-speed genome organization where specific genomic regions are characterized by extensive presence-absence polymorphisms between strains (Ma *et al*., 2010; van Dam *et al*., 2017; Fokkens *et al*., 2018; Zhang *et al*., 2020). These variable accessory regions (ARs) can be embedded in core chromosomes or can encompass entire accessory chromosomes (Ma *et al*., 2010; Fokkens *et al*., 2018). Interestingly, shared ARs in otherwise genetically diverse strains have been linked to the adaptation towards the same host (Fokkens *et al*., 2018; Henry *et al*., 2021). Moreover, ARs can transfer the capacity to infect a specific host between isolates (Ma *et al*., 2010; Ayukawa *et al*., 2021). For example, transfer of accessory chromosome 14 from a tomato infecting *Fusarium* (*Fol*4287) to a non-pathogenic *Fusarium* strain converts the non-pathogenic isolate into a tomato infecting pathogen (Ma *et al*., 2010). This transfer of pathogenicity is likely linked to effector genes that are localized in ARs such as the 14 well-studied *Secreted In Xylem* (*SIX*) effectors (Rep *et al*., 2002). Consequently, the presence and absence of effectors can group genetically diverse *F. oxysporum* strains based on their host range (van Dam *et al*., 2016). Whole-genome sequencing of diverse *F. oxysporum* can provide novel insights into the origin of host specificity (Fokkens *et al*., 2018; Henry *et al*., 2021). A recent population study of *F. oxysporum* isolates infecting chickpea suggests that, contrary to expectations (Gordon & Martyn, 1997; Henry *et al*., 2021), *F. oxysporum* might undergo sexual as well as clonal reproduction (Fayyaz *et al*., 2023), which could impact the constitution and the dynamics of ARs and, ultimately, influence pathogenicity. However, despite the importance of ARs for host adaption, little is known about the constitution of ARs in *F. oxysporum* strains that cause disease in the same host, and the origin and processes that shape the evolution of ARs in *Fusarium*.

Fusarium wilt of banana (FWB), which is caused by a suite of *F. oxysporum* species (Maryani *et al*., 2019), is a major constraint for banana production and threatens food security in tropical and subtropical countries where >400 million people depend on banana cultivation (FAO, 2020). In *F. oxysporum* species causing FWB, three different races can be distinguished based on their pathogenicity towards subsets of banana varieties; Race 1 (R1) strains cause FWB in Gros Michel, Race 2 (R2) strains infect Bluggoe, and Tropical Race 4 (TR4) strains infect Gros Michel, Bluggoe, and Cavendish (Ploetz, 2015). In addition to TR4, subtropical race 4 (STR4) strains can cause disease in Cavendish banana under environmental stress conditions, for example caused by lower temperatures in subtropical regions (Ploetz, 2006). All races infect additional locally important banana varieties. TR4 has developed into a pandemic over the last 60 years (Ordonez *et al*., 2015; van Westerhoven *et al*., 2022b) and is of particular concern as it causes FWB in Cavendish, the banana variety that dominates global productions (>50%) and export trade (>95%) (FAO, 2020). Despite the impact of FWB on banana production, we know little about the ARs in *Fusarium* strains that infecting banana and their relation to pathogenicity.

Here we analyze the dynamics of ARs in a suite of *Fusarium* strains causing FWB. To gain insights into the genomic structure, we generate seven chromosome-level reference genome assemblies for R1, R2 and TR4 strains, and use a pan-genomic framework together with a global panel of 69 strains to gain insights into the evolution of ARs in genetically diverse *Fusarium* species.

## Materials and Methods

### Collection of *Fusarium* strains infecting banana

We analyzed a diverse set of 69 *Fusarium* strains infecting banana (**Supplementary Table 1**). *Fusarium* strains were isolated from different banana growing regions worldwide directly from the pseudostem of symptomatic banana plants. Here, 35 *Fusarium* strains were newly sequenced and we included 34 publicly available whole-genome sequencing datasets (based on short-read sequencing approaches) (Guo *et al*., 2014; García-Bastidas *et al*., 2014; Zheng *et al*., 2018; Warmington *et al*., 2019; Maymon *et al*., 2020; Garcia-Bastidas *et al*., 2020; van Westerhoven *et al*., 2022a). To obtain high-quality reference assemblies, seven strains were sequenced with Oxford Nanopore Technology (ONT).

### Whole-genome sequencing

*Fusarium* isolates were cultured on potato dextrose agar (PDA) and conidia were transferred to liquid media for incubation at 25°C. Fungal material was isolated from liquid culture for genomic DNA extraction. The exact DNA isolation protocol per strain can be found in Supplementary Methods. To obtain long-read, whole-genome sequencing data, seven strain (36102, II5, CR1.1, C058, C135, C081, and M1) were sequenced using Oxford Nanopore Technologies (Oxford, UK). An R9.4.1 flow cell was loaded and run for 24h. Base calling was performed using Guppy (version 3.1.5; C058, C135, C081, and M1) and MiniKNOW (version 3.4.6; II5, 36102, and CR1.1). The other *Fusarium* strains were sequenced by Beijing Genomic Institute (BGI) using Illumina paired-end sequencing, 31 were sequenced during this study and 38 were obtained during other studies (Guo *et al*., 2014; Ordonez *et al*., 2015; Zheng *et al*., 2018; Warmington *et al*., 2019; Maymon *et al*., 2020; Garcia-Bastidas *et al*., 2020; van Westerhoven *et al*., 2022a).

### Gene expression analysis

Gene expression patterns were compared between *in vitro* growth and 8 days post infection. Cavendish ‘Grand Naine’ banana plants were grown in a greenhouse compartment (28 ± 2^◦^C, 16 h light, and ∼85% relativity humidity) and acclimatized under plastic for 2 weeks to maintain high humidity. The roots of ∼2.5 months old plants were dip inoculated with 10^6^ spores per ml. RNA was isolated from the roots of inoculated plants at 8 days post inoculation and from mycelium grown on PDA medium. Samples were ground and RNA extraction was performed using the Maxwell Plant RNA kit (Promega) following manufacturer’s instructions. The quality of the RNA samples was tested by agarose electrophoresis and quantified by Nanodrop (ThermoFisher), and subsequently sent to KeyGene (The Netherlands) for sequencing. Quantifications of gene transcripts from sequenced RNA-seq reads was performed using Kallisto (Bray *et al*., 2016).

### Genome assembly and annotation

We obtained chromosome-level *de novo* genome assemblies for seven Fusarium isolates sequenced by ONT long-read technology. Adapter sequences were removed with Porechop (version 0.2.4, default settings; https://github.com/rrwick/Porechop/tree/master) and reads were self-corrected, trimmed, and assembled using Canu (Koren *et al*., 2017) (version 1.8). Genome assemblies were polished using ONT raw reads with Racon (Vaser *et al*., 2017) (version 1.5.0) and with high-quality Illumina short read using two rounds of Pilon(Walker *et al*., 2014) (version 1.24). Contigs were scaffolded, if needed, using longstitch (Coombe *et al*., 2021) (version 1.0.2).

Fusarium strains sequenced solely with short-read sequencing technology were *de novo* assembled with Spades (Bankevich *et al*., 2012) (version 3.13.0, with default settings). Contigs shorter than 500 bp were subsequently removed. To assess the assembly quality, the genome assemblies were analyzed with QUAST (Gurevich *et al*., 2013) (version 5.0.2), and genome completeness was estimated based on the presence of conserved single-copy genes using BUSCO (Simão *et al*., 2015) (version 5.1.2) with the *hypocreales* odb10 database.

We performed gene annotation of 69 *Fusarium* genome assemblies using Funannotate (Palmer & Stajich, 2019) (version 1.8.9) with the *Fusarium oxysporum* f. sp. *lycopersici* 4287 Uniprot proteins, uniport database, and previously predicted effectors (van Dam *et al*., 2016) as external evidence. Genomes were masked using RepeatMasker (version 4.1.1) following repeat prediction with RepeatModeler (http://www.repeatmasker.org). For *Fusarium* strains II5, CR1.1, and 36102, RNA-seq data was used to guide gene model prediction, and *ab initio* training parameters estimated using RNA-seq data of II5 were re-used to support the annotation in other genome assemblies. Effectors genes were predicted based on their association with <u>M</u>iniature <u>IMP</u>ala (MIMP) elements in *Fusarium* (van Dam *et al*., 2016). Briefly, ORFs were predicted 5 kb up- and downstream to MIMPs (‘AGT[GA][GA]G[GAT][TGC]GCAA[TAG]AA’) and these were refined using AUGUSTUS (version 3.3.3)(Stanke & Morgenstern, 2005), with the *Fusarium* species specified. In the set of predicted MIMP-genes, secreted proteins were predicted using SignalP (version 5) and subsequently effectors were predicted using EffectorP (version 3.0). The predicted effectors were added to the Funannotate annotation using AGAT (version 0.8.0.)

To recover potentially missing genes in the gene annotations, we used the comparative annotation toolkit (Fiddes *et al*., 2018) (version 2). Cactus (Armstrong *et al*., 2020) (version 2.0.1) was used to create whole-genome alignments of all 69 genomes, using the Minigraph-Cactus approach (Hickey *et al*., 2023). Consecutively, the CAT was used to carry-over gene annotations based on Funannotate gene predictions with the RNA-seq data (with Augustus settings pre trained on II5, --global-near-best 1, --augustus-utr-off, --augustus-cgp, --maf-chunksize 550000 --maf-overlap 50000), which added approximately 200 genes per genome assembly.

### Core gene phylogeny

The phylogenetic relationship between *Fusarium* strains infecting banana together with 55 publicly available non-banana infecting *Fusarium* strains was inferred from 3,811 conserved single-copy BUSCO genes. Protein sequences were aligned using Mafft (Katoh *et al*., 2002), (version 7.453) and a maximum-likelihood phylogeny was determine using IQtree (Chernomor *et al*., 2016) (version 1.6; -m MFP+MERGE, -B 1000 with 1,000 ultra-fast bootstrap), *Fusarium verticillioides* (ASM14955v1 (Cuomo *et al*., 2007)) was included as an outgroup.

### Pangenome analysis

We detected conserved and accessory regions in the set of *Fusarium* isolates by performing all-vs-all whole-genome alignments with nucmer (Kurtz *et al*., 2004) (version 3.1; --max-match). We removed small alignments with delta-filter (-l 5000) and only retained the best match per alignment (-1). Genomic regions present in all 69 *Fusarium* isolates were considered core, regions covered by less than 80% were considered accessory, and regions between these values were considered softcore. Pairwise similarity between accessory regions was calculated using custom python scripts (https://github.com/Anouk-vw/Fusarium_pg) and alignments were visualized with PyGenomeViz (https://github.com/moshi4/pyGenomeViz) and Circlize (Gu *et al*., 2014).

To identify orthologous groups, we used Broccoli (Derelle *et al*., 2020) (version 1.1) on the predicted protein-coding genes from the 69 *Fusarium* strains infecting banana; per gene, only the longest transcript was included. OGs present in all isolates were considered core, OGs found in more than 55 genomes (80%) were considered softcore, and genes found in less than 55 of the genomes were considered accessory.

We determined the number of non-synonymous and synonymous substitutions (dN/dS) per OG. All genes in an OG were aligned using mafft (Katoh *et al*., 2002) (version 7.490) and non-synonymous and synonymous substitutions per gene pair in the OG were inferred based on a codon-guided nucleotide alignment, created by pal2nal (Suyama *et al*., 2006) (version 14), with codeML (from paml (Yang, 2007), version 4.9). Differences between groups of genes were further analyzed using custom python scripts and compared using a two-sided Mann-Whitney U test in SciPy (Virtanen *et al*., 2020) (version 1.10.1).

### Estimation of gene age

Orthofinder (Emms & Kelly, 2019) (version 2.5.4) was used to detect the presence of homologs in 308 phylogenetically distinct fungal species from the joint genome institute; per fungal family, one fungal genome assembly was randomly selected (Supplementary table **5**). Homologs that occur at least once outside the phylum Ascomycota are considered *old* homologs, homologs occurring only within Ascomycota are further subdivided into *Hypocreales* and *Fusarium*.

### Detection of duplication events

We detected gene duplication events by self-comparisons of the translated protein sequences (only the longest isoform) of the chromosome-level genome assemblies of *Fusarium* infecting banana, as well as *Fusarium oxysporum* f. sp. *lycopersici* (strain *Fol*4287 (Ma *et al*., 2010); GCF_000149955.1, *Fusarium* graminearum (Cuomo *et al*., 2007) (GCF_000240135.3), *Fusarium verticillioides* (Cuomo *et al*., 2007) (GCF_000149555.1), and *Fusarium solani* (Mesny *et al*., 2021) (GCF_020744495.1). Per *Fusarium* strain, the predicted protein sequences were compared with BLAST and the best five hits were retained after filtering for an e-value 1e^-10^ and query coverage larger than 50%. Then, BLAST alignments were used to detect collinear duplicated blocks with MCscanX (Wang *et al*., 2012); blocks of five genes were considered collinear and matches were filtered with the parameters match-score 50 and e-value 10^-5^. Subsequently, we classified duplicated genes into dispersed, proximal, tandem, or segmental duplications with MCscanX’s ‘duplicate_gene_classifier’ script.

Copy number variations were identified by mapping short-read, whole-genome sequencing datasets against the chromosome-level genome assembly of *Fusarium* strain II5 using BWA-mem (Li, 2013) (version 0.7.17). We determined the sequencing coverage over the reference genome with bedtools (version 2.30), which was visualized Wgscoverageplotter from jvarkit (Lindenbaum, 2015) (version 1.1.0), filtering reads with a mapping quality of 0. Single nucleotide polymorphisms were called using GATK (version 4.0.2; -ploidy 1) and filtered according to GATK best practices (Van der Auwera *et al*., 2013).

## Results

### Fusarium strains infecting banana are genetically diverse

To study the occurrence and evolution of ARs in *Fusarium* strains infecting banana, we sequenced and assembled 69 strains (Fig. **1a**, Supplementary table **1**) that were isolated from all major banana growing regions and classified into R1 (39 strains), R2 (two strains) and TR4 (28 strains) (Fig. **1a,b**, Supplementary table **1**). *De novo* assembly of seven long-read sequencing strains infecting different banana varieties (three TR4, two R1, two R2) achieved chromosome-level contiguity containing 12 to 15 contigs, most of which represent complete chromosomes flanked by telomeric repeats (Supplementary table **2,3**). While genome assemblies obtained solely from short-read data yielded more fragmented assemblies, all assemblies contained at least 96.9% of the single-copy BUSCO genes. The assemblies range from 43 to 51 Mb in size (Fig. **1b**) and encode between 15,664 and 17,865 predicted proteins, of which 551 to 671 are effector candidates, and consist for 2.6 – 8.9% of repeats (Fig. **1e,f**), similar to previous *Fusarium* genome assemblies (van Dam *et al*., 2016; Warmington *et al*., 2019; Zhang *et al*., 2020).

**Figure 1.**
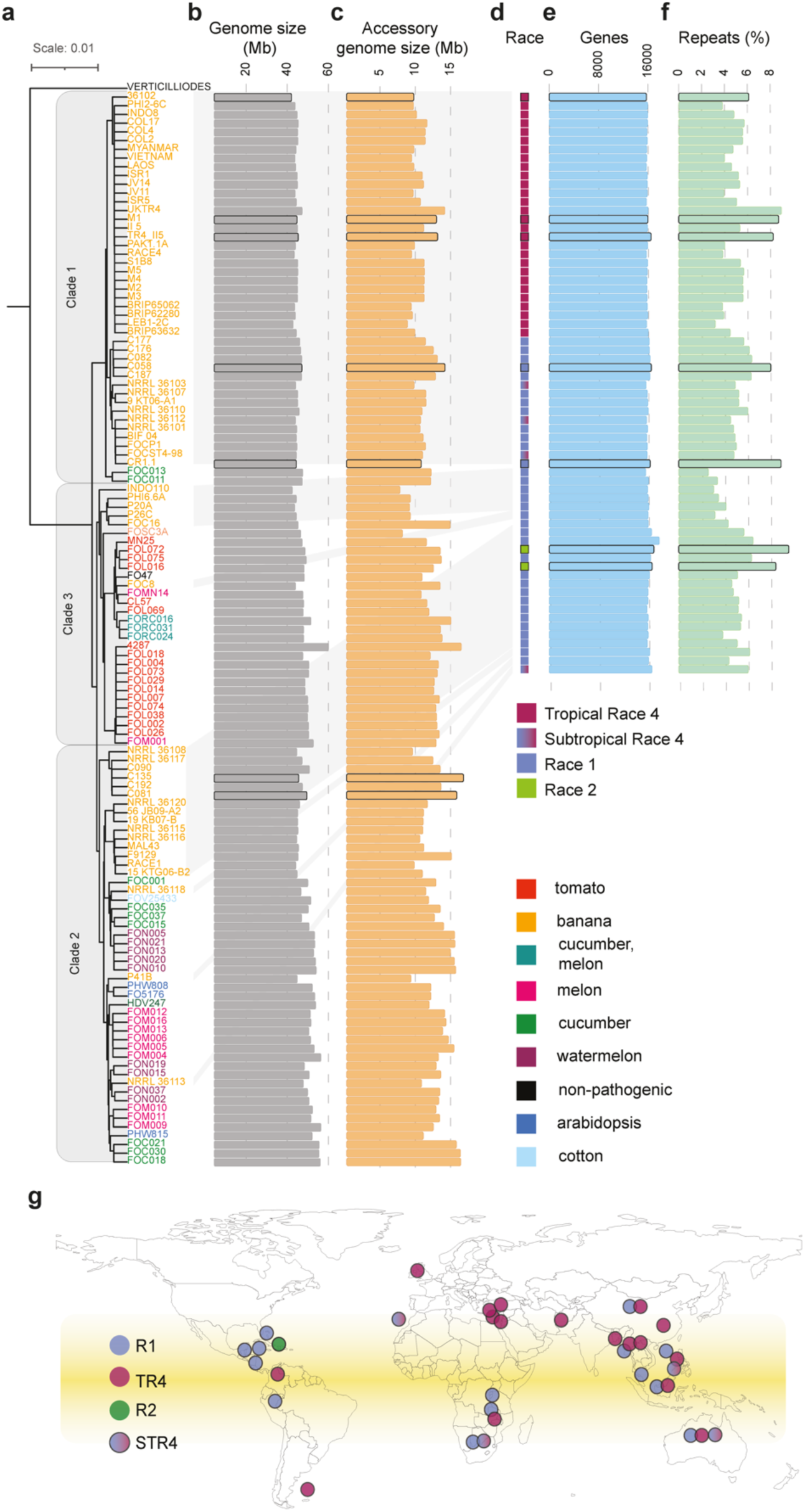
*Fusarium* strains infecting banana are genetically diverse. **a**) *Fusarium* strains that infect banana are polyphyletic (yellow labels). The relationship between *Fusarium* infecting banana and *Fusarium* strains that infect other plant hosts (see label color) is estimated using maximum-likelihood phylogenetic analysis based on 3,811 concatenated conserved single-copy BUSCO genes. Black outline indicates high quality genome assemblies. **b)** Genome sizes of *Fusarium* strains range between 43 to 61 Mb. **c**) The sizes of accessory regions in *Fusarium* strains range from 6.7 to 16.5 Mb. **d**) TR4 strains (purple squares) occur in clade 1 and are genetically similar, as well as the two R2 strains (green). By contrast, the R1 strains (blue squares) are polyphyletic. **e**) *Fusarium* strains that infect banana encode between 15,664 and 17,865 genes and **f**) between 2.9 and 8.9% repeats. **g)** The collection of 69 *Fusarium* strains originates from different banana growing regions worldwide; the approximate origin of isolation is indicated by dots, colored according to the race designation (R1, R2, STR4, and TR4). Strains sequenced by long-read sequencing technology are highlighted in black.

To assess the genetic diversity of *Fusarium* strains infecting banana in relation to 55 publicly available *Fusarium* strains infecting eight different host plants (van Dam *et al*., 2016) (Supplementary table **1**), we conducted a phylogenetic analysis based on 3,811 single-copy orthologous genes (2,226,902 amino acid positions). As expected, *Fusarium* strains infecting banana are polyphyletic and associated with *F. oxysporum* clades 1, 2, and 3 (Fig. **1a**)(O’Donnell *et al*., 1998; Maryani *et al*., 2019). We observed low nucleotide diversity between TR4 strains (pi-nucleotide diversity 0.0018) compared to a 100x higher diversity between R1 strains (pi-nucleotide diversity 0.185), corroborating that TR4 strains evolved from a single recent clonal origin (Ordonez *et al*., 2015; Maryani *et al*., 2019). Based on their genetic diversity, *Fusarium* strains infecting banana have recently been separated into different species, and the lineage that encompasses TR4 strains (Fig. **1d**) is now referred to as *Fusarium odoratissimum* (Ordonez *et al*., 2015; Maryani *et al*., 2019).

### *Fusarium* strains causing Fusarium wilt of banana have a compartmentalized genome

To identify ARs in *Fusarium* strains infecting banana, we performed whole-genome alignments of the seven chromosome-level genome assemblies to the reference genome assembly of the tomato infecting *F. oxysporum* strain 4287 (*Fol*4287)(Ma *et al*., 2010). We observed eleven homologous chromosomes between the TR4 reference strain II5 and *Fol*4287, that are considered the core genome (chromosome 1-11; Fig. **2a**). In addition to the core chromosomes, we observed two large ARs specific for II5. The first spans 1.8 Mb occupying 27% of chromosome 1 in proximity to one telomere and the second region is 1.1 Mb in size and constitutes the entire contig 12 (Fig. **2a**). Contig 12 only contains one telomere (Fig. **2b**), and therefore it remains unknown whether this accessory region represents an extra chromosome or is attached to one of the core chromosomes. Importantly, we assemble a similar contig in TR4 strain M1, and a similar contig is present in the independently assembled TR4 strain Eden (Warmington *et al*., 2019), which strongly suggests that this contig represents an separate accessory region.

**Figure 2.**
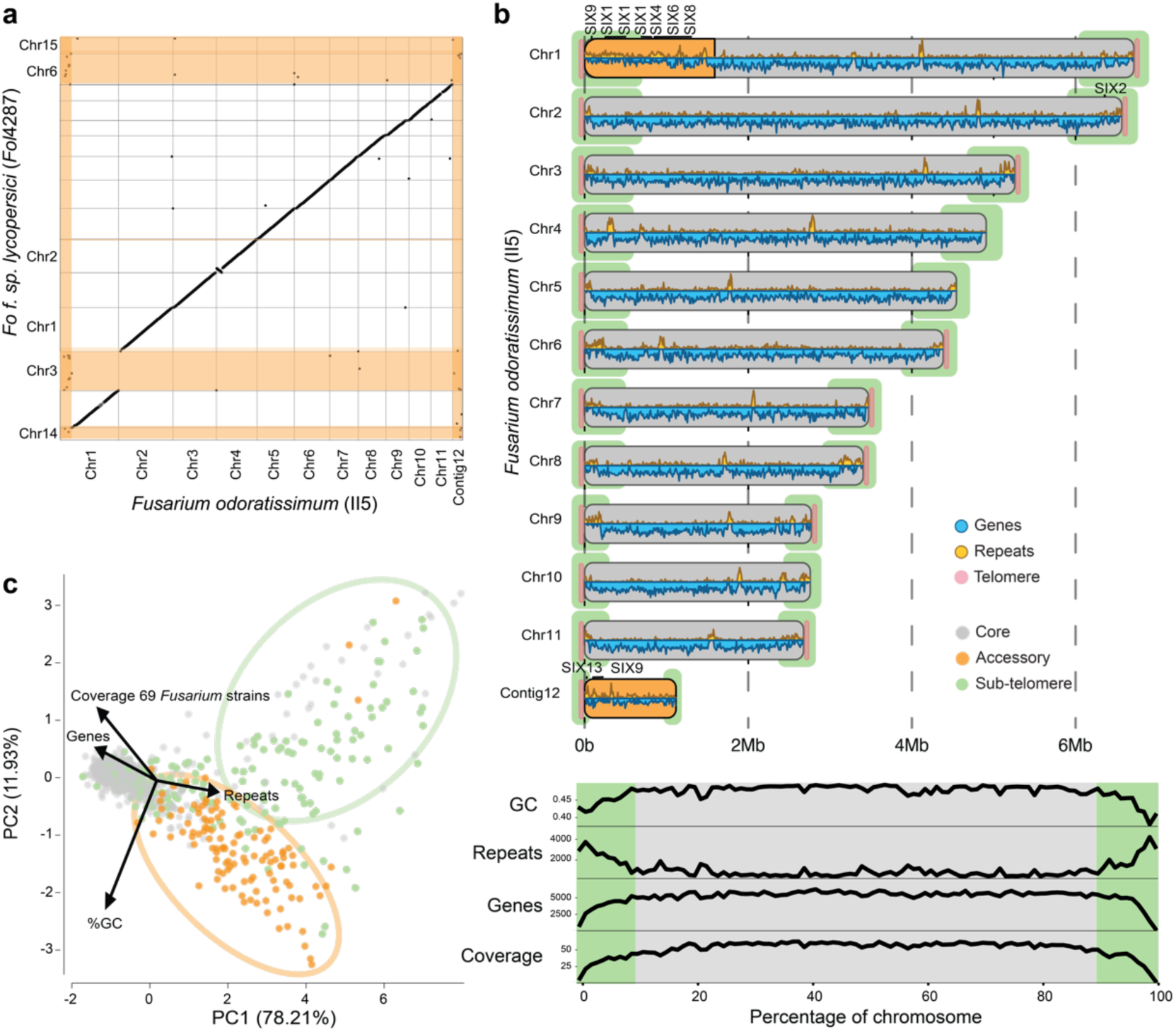
*Fusarium* strains infecting banana carry accessory regions (ARs). **a)** The TR4 reference strain II5 contains eleven core chromosomes (chromosome 1-11) as well as two ARs (1,8 Mb on chromosome 1 and entire contig 12). Core chromosomes align to *Fol*4287 in contrast to the ARs that are highlighted in orange. **b**) The ARs (orange) in II5 carry most *Secreted In Xylem* (*SIX*) genes and show an increased repeat and lower gene content. A similar pattern is found in the sub-telomeric regions (green, first and last 10% of the chromosome). **c**) Principal component analysis on gene-, repeat-, GC-content, and coverage of 69 *Fusarium* strains separates the genomic regions of II5 into three different clusters. ARs (orange and green) are separated from the core region (grey) and further separated into those ARs localized in sub-telomeric regions (green) and those on chromosome 1 and contig 12 (orange).

Compared to the eleven core chromosomes, the two ARs have an increased repeat content (28.30% vs 6.41%) and a decreased gene density (32.15% vs 59.71%) and GC content (46.4% vs 47.5%) (Fig. **2b**). Importantly, the ARs in II5 encode 463 genes, nine are homologous to *SIX* genes (Fig. **2b**) and 22 are predicted effectors (out of 629 predicted effectors). Although this suggests that the ARs are important for pathogenicity on banana, they are not collinear with other *Fusarium* strains infecting banana (Supplementary Fig. 1).

To further analyze the diversity of ARs, we identified ARs in our global collection using a pan-genomic approach based on all-vs-all whole genome alignments. ARs were defined as regions longer than 5 kb that are absent in more than 55 of 69 strains (80%). We identified a varying amount of ARs in *Fusarium* strains infecting banana (Fig. **1c**), ranging from 6.7 Mb (15% of the genome size) in strain Indo110 (R1) to 15.5 Mb (30%) in strain C135 (R2), in line with the 19 Mb (29%) of ARs in the reference strain *Fol*4287 (Ma *et al*., 2010). Next to the two ARs in chromosomes 1 and contig 12, our pan-genomic approach identified additional 8 Mb of ARs in II5, typically localized at sub-telomeric regions (defined as the first and last 10% of each chromosome) (Fig. **2b**). These sub-telomeric regions contain 1,287 genes, of which 80 are predicted effector genes (6.6%). We also identified that chromosomes 9 (42.6% ARs, 87 SNPs per kb), 10 (27% ARs, 80 SNPs per kb), and 11 (36% ARs, 92 SNPs per kb) are less conserved than the other core chromosomes (on average 20% ARs, 59 SNPs per kb) (Supplementary Fig. 2), which resembles the situation of the previously identified ‘fast-core’ chromosomes in *Fol*4287 (Fokkens *et al*., 2018).

We show that ARs can be defined based on sequence conservation and differ from the core genome in gene, repeat and GC-content. Consequently, principal component analyses distinguished ARs from the core genome based on these four features (Fig. **2c**). Unanticipatedly, this analysis also separated the large ARs on chromosome 1 and contig 12 from the ARs localized at the sub-telomeres (Fig. **2c**), this separation is driven by the slightly lower GC content in sub-telomeric regions (44% vs 47%). A reduced GC-content can be caused by repeat induced polymorphisms (RIP), a fungal defense mechanisms that introduces C to T mutations in repetitive regions (Galagan & Selker, 2004). In the sub-telomeric region, 25.4% of the repetitive elements are likely subjected to RIP (i.e., composite RIP index > 0), in contrast to only 7.7% in the other ARs, suggesting that repetitive elements at the sub-telomeres are older, or alternatively, that RIP has been more active.

### Highly variable accessory regions in *Fusarium* strains infecting banana

To characterize the similarity of ARs in *Fusarium* strains infecting banana, we compared the ARs from the seven chromosome-level genome assemblies. Notably, the ARs are highly diverse with little sequence and structural similarity (Fig. **3**, Supplementary Fig. 1). In TR4, the ARs from strain II5 on chromosomes 1 and contig 12 show extensive similarity to the corresponding chromosomes in strain M1 (Fig. **3a**, Supplementary Fig. 1). On the other hand, isolate 36102, causing milder symptoms in Cavendish than TR4 II5, carries only the AGR on chromosome 1, and does not encode a region similar to contig 12 (Supplementary Figure 1). None of the AGRs from strain II5 are present in the strains representing Race 1 and Race 2. Similar to the TR4 strains, the accessory contigs of two Race 2 strains correspond to each other (Supplementary Fig. 1), but no shared ARs are found among R1 strains. Moreover, we observed that no ARs are conserved among different races, for example only limited genetic material is shared between ARs of TR4 and R2 (Supplementary Fig. 1). The diversity of ARs was further corroborated by pairwise analyses of all 69-strains, which revealed that, on average, TR4 shared 91% of the ARs, whereas R1 shared only 24% (Fig. **3b**). In line with the high number of shared ARs in TR4 (*F. odoratissimum*, Fig. **3b**), the amount of shared ARs was highest among R1 strains belonging to the same *Fusarium* species (60% shared ARs, Fig. **3b**, Supplementary Fig. 3), however R1 strains from different species share very few ARs (Fig. **3b**).

**Figure 3.**
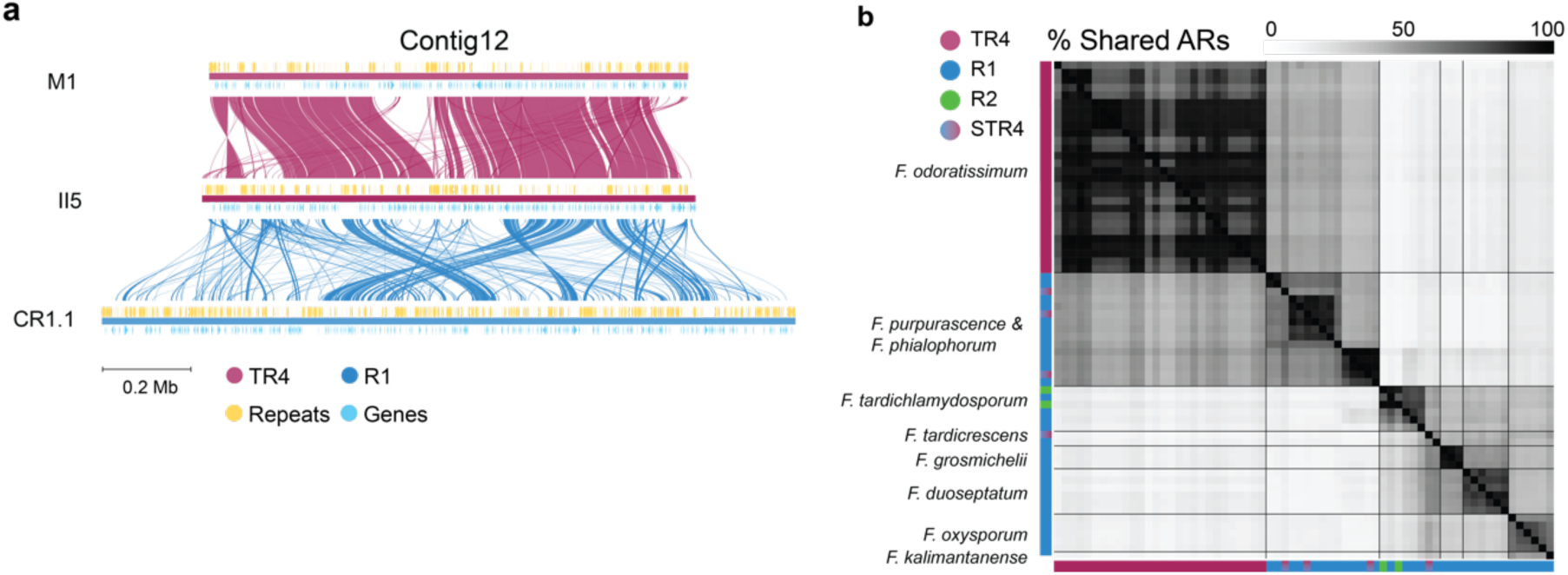
Accessory regions in *Fusarium* strains infecting banana differ significantly. **a**) Accessory contig 12 of the TR4 reference strain II5 is shared with TR4 strain M1, however this contig shows little similarity to R1 strain CR1.1. Lines (red and blue) indicate aligned regions between strains. Small alignments are present between R1 and TR4 (blue lines), however these are rearranged and often involve repetitive sequences (yellow blocks). **b**) R1 strains (39 strains, highlighted in blue) have a diverse accessory content sharing on average only 20% of the ARs, whereas TR4 strains (28, highlighted in purple) share most of their ARs (91%). Phylogenetically related strains, here indicated by their species name (Maryani *et al*., 2019), share more ARs than distantly related strains.

To quantify to which extent *Fusarium* strains infecting banana share ARs with *Fusarium* strains infecting different plant hosts, we identified and compared ARs in a set of 55 additional *Fusarium* strains that are pathogenic to different plant hosts (van Dam *et al*., 2016). We observed that genetically related strains share most ARs (Supplementary Fig. 4), however we also noticed that *Fusarium* strains infecting banana encode more diverse ARs than for instance the tomato-infecting strains; on average, *Fol* strains share 62% ARs, with a minimal 30% of the ARs shared between any two *Fol* strains (Supplementary Fig. 4). In contrast, banana-infecting strains on average share 44% of the ARs and a pair of strains can share as little as 12% (Supplementary Fig. 4). Thus, we demonstrate that ARs are highly variable among *Fusarium* strains infecting banana and, importantly, strains do not share one host-associated chromosome in contrast to what was previously reported for *Fusarium* infecting tomato (Fokkens *et al*., 2018).

### Evolutionary dynamics of the accessory genome

*Fusarium* strains infecting banana do not share large ARs (Fig. **3b**), but we reasoned that ARs nevertheless might share genes that contribute to the pathogenicity towards banana. We therefore grouped 1.1 million protein-coding genes predicted in 69 strains into 22,612 orthologous groups, and subsequently analyzed the gene-content diversity. The pan-genome consisted of 12,101 core groups (53%) that are present in all 69 strains, 2,395 softcore groups present in at least 80% of the strains, 5,595 (25%) accessory groups present in less than 80% of the strains, and 2,521 unique genes (Fig. **4a**). Importantly, the pan-genome based on our collection captures the diversity of protein-coding genes in *Fusarium* strains infecting banana as we did not observe an increase in the pan-genome size after including more than 40 strains (Fig. **4a**). The conserved core genes are, as expected, enriched with genes with housekeeping functions, whilst accessory genes are enriched with genes encoding secondary metabolites (Supplementary Fig. 5). No enrichment of effector genes is found in any of the three categories (core, accessory, or softcore); the pan-genome consists of 739 gene families encoding predicted effectors that are evenly distributed over the different gene categories (3.7% of accessory genes and 3.5% of the core genes). The role of accessory genes in host infection is suggested by gene expression, as accessory genes are upregulated eight days after banana infection, with a median log2fold change of 1.59 (compared with the *in vitro* control) (Fig. **4b**), comparable to the upregulation of effector genes (median log2fold change of 2.4). The core genes, however, show significantly less increase of gene expression upon plant infection (median log2fold change = 0.25, P < 0.05, two-sided Mann-Whitney U test) (Fig. **4b**). Although this suggests a role of accessory genes in host-pathogenicity, we observed a highly diverse gene content in R1 strains with only 246 out of 7,832 accessory genes (3%) being shared among all R1 strains (Fig. **4f**). TR4 strains, in contrast, have a more similar gene profile and share 2,493 of the 4,406 accessory genes (57%).

**Figure 4.**
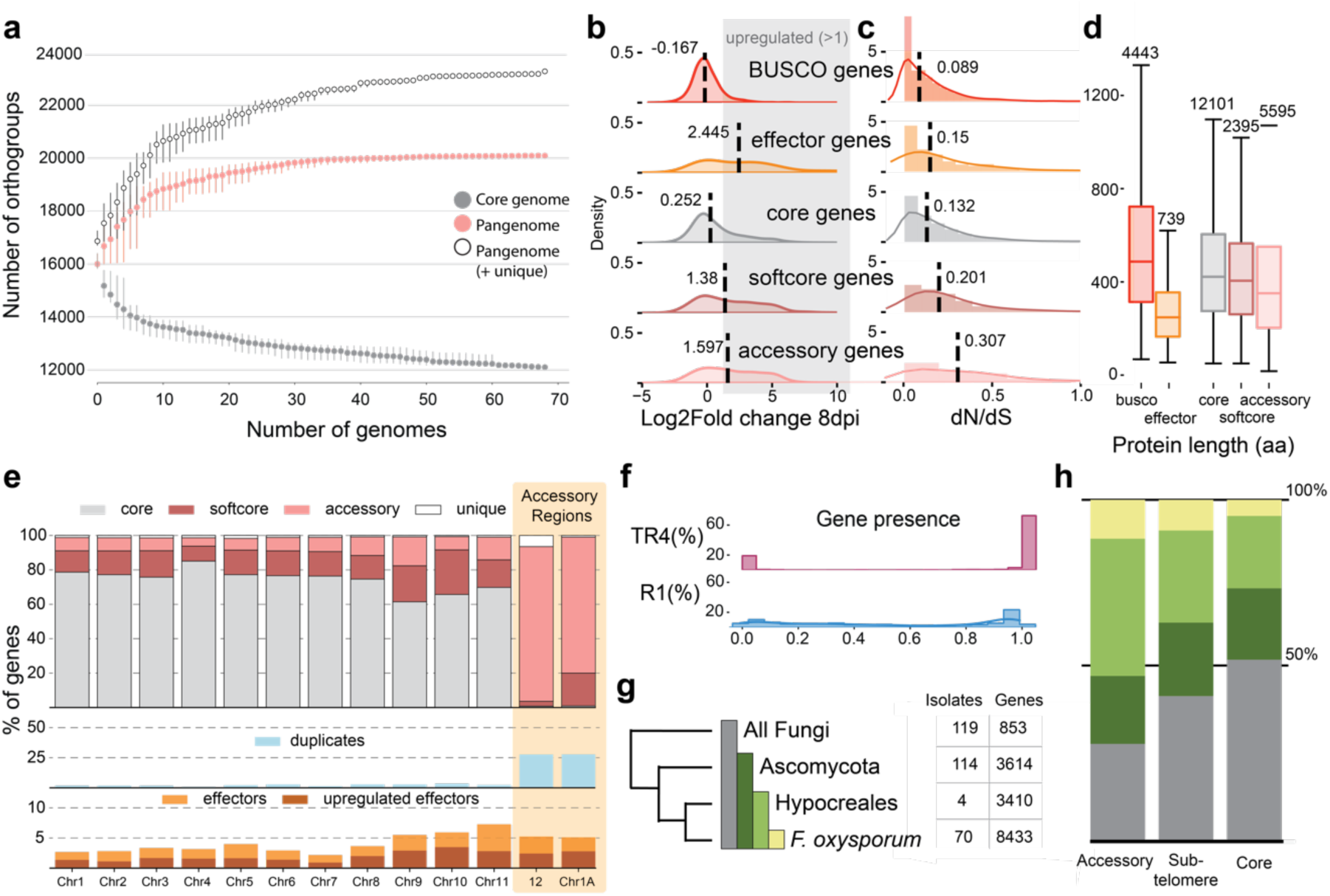
Evolutionary dynamics of genes in the different genomic compartments in *Fusarium* strains infecting banana. **a)** Global collection of *Fusarium* infecting banana has a pan-sgenome size of 20,091 orthologous groups (OG, 22,612 when unique genes are included), of which 12,101 are present in all strains (grey). The pan-genome is saturated after adding 40 genomes. The order of genomes was randomly sampled ten times, dots show the average pangenome size and the vertical grey lines indicate the standard deviation. **b**) Accessory genes are upregulated *in planta* at 8 days post inoculation of banana plants. Average log2fold change of accessory genes (2,416 in II5) and effector genes (629 in II5) shows these genes are typically upregulated upon plant infection, which contrasts the core (11,740 genes, log2-fold change = 0.252) and BUSCO genes (4,364 genes, log2-fold change = -0.167). The median log2fold change of all gene categories is significantly different (p<0.05, two sided Mann-Whitney U test). **c**) Sequence conservation estimated by dN/dS values. BUSCO gene families (4,443 OGs) and core gene families are most conserved, whereas the accessory genes evolve under more relaxed selective pressure. All gene categories show significant variation in dN/dS value (p<0.05, Mann-Whitney U test). **d)** Distribution of protein length separated by different gene categories shows that accessory gene families (5,595 OGs, pink) are shorter (median length of 315 aa per OG) than core gene families (median length 469 aa per OG, 12,101 OGs, grey). All gene categories differed significantly from each other (p<0.05, Mann-Whitney U test). **e**) ARs in II5 (orange, contig 12 and chromosome 1A) consist mostly of accessory genes (pink, 405 out of 463, 87%) and 28% of these genes are part of expanded orthogroups (129 out of 463). Accessory genes can also be found in core chromosomes. **f**) Gene content is variable between R1 strains, and only 20% of the genes are present in 80-90% of the R1 strains. Most genes (60%) in TR4 are present in all TR4 strains (90-100%). **g)** Gene families in TR4 strain II5 differ to which extend they have homologs in 308 fungal phyla; most genes are present in related *Fusarium* strains, but only a few gene families (853) are conserved among all considered fungi. **h**) Accessory regions contain most recent genes, present in *Fusarium* (51) and Hypocreales (160). Genes in core regions are evolutionarily older with more genes that have homologs in ‘all fungi’ (7,764 out of 11,740 genes).

In TR4 strain II5, ARs have a different gene composition than the core regions, with ARs consisting mostly of accessory genes (405 out of 463; 87%), however four core and 38 soft core genes are also found in accessory regions (Fig. **4e**), indicating that some shared genes can be found between otherwise non-conserved regions. ARs seem to contain recently evolved genes, as most AR genes (71%) have homologs in closely related fungi (Fig. **4g,h**) and only 29% are considered “old”, i.e., with homologs in all fungi, which contrasts to the 53% old genes in the core genome (7,764, out of 14,774 genes) (Fig. **4g,h**). Additionally, accessory genes are significantly shorter (315 aa) and evolve under relaxed selective pressure (dN/dS = 0.20), compared to the core genes (469 aa, dN/dS = 0.089; P < 0.05, two-sided Mann-Whitney U test) (Fig. **4b,c**), both hallmarks of more recent genes (Wolf *et al*., 2009). This suggests that most genes in the ARs evolved recently, and that these regions serve as cradles for the evolution of novel genes. In addition, many genes on the ARs (177 out of 463) belong to expanded gene families based on the defined orthogroups (Fig. **4e**), suggesting that gene duplications play a role in the evolution of ARs in *Fusarium* strains infecting banana.

### Variable effector repertoire in Fusarium oxysporum genomes

Effector gene repertoires define host range (van Dam *et al*., 2016; Brenes Guallar *et al*., 2022), and we anticipated that effector presence-absence profiles would distinguish *Fusarium* strains infecting banana from *Fusarium* strains infecting other hosts. We predicted candidate effector genes in all 124 *Fusarium* isolates based on their proximity to Miniature IMPala elements (MIMP) that are known to co-localize with some effector genes in *F. oxysporum* (van Dam *et al*., 2016; Brenes Guallar *et al*., 2022), which yielded 398 MIMP-associated effector gene families (160 of these were predicted in the complete set of 739 candidate effectors). The predicted MIMP-associated effector repertoires clustered *F. oxysporum* isolates on host range (van Dam *et al*., 2016; Brenes Guallar *et al*., 2022) (Fig. **5**). While banana-infecting strains form a clear cluster, only four MIMP-associated effectors were identified that occur in most (i.e., >80%) of banana-infecting strains but are consistently absent (i.e., present in less than 20%) in other *F. oxysporum* strains. Moreover, races within *Fusarium* strains infecting banana are not clearly separated; TR4 strains have very similar effector profiles, yet R1 strains differ considerably and do not encode shared effectors. Interestingly, two STR4 strains carry an effector profile remarkably similar to the effector profile of TR4 strains (Figure 5). This suggests that this effector profile contributes to the pathogenicity of these STR4 strains to Cavendish banana. However, this effector profile is not STR4 specific as the other two STR4 strains encode effector profiles similar to R1 strains (Figure 5). This difference can be the result of misclassification of STR4 strains, because STR4 infects Cavendish only under environmental stress conditions and this environmental component makes it difficult to distinguish true STR4 isolates from Race 1 isolates (Ploetz, 2006). To further investigate effectors shared between TR4 and STR4, an accurate screening strategy to distinguish true STR4 needs to be conducted. We here reveal a diverse effector repertoire in R1 strains, together with the variable ARs and gene content, this suggests that the species encompassing R1 utilize varying molecular mechanisms to support banana infection.

**Figure 5.**
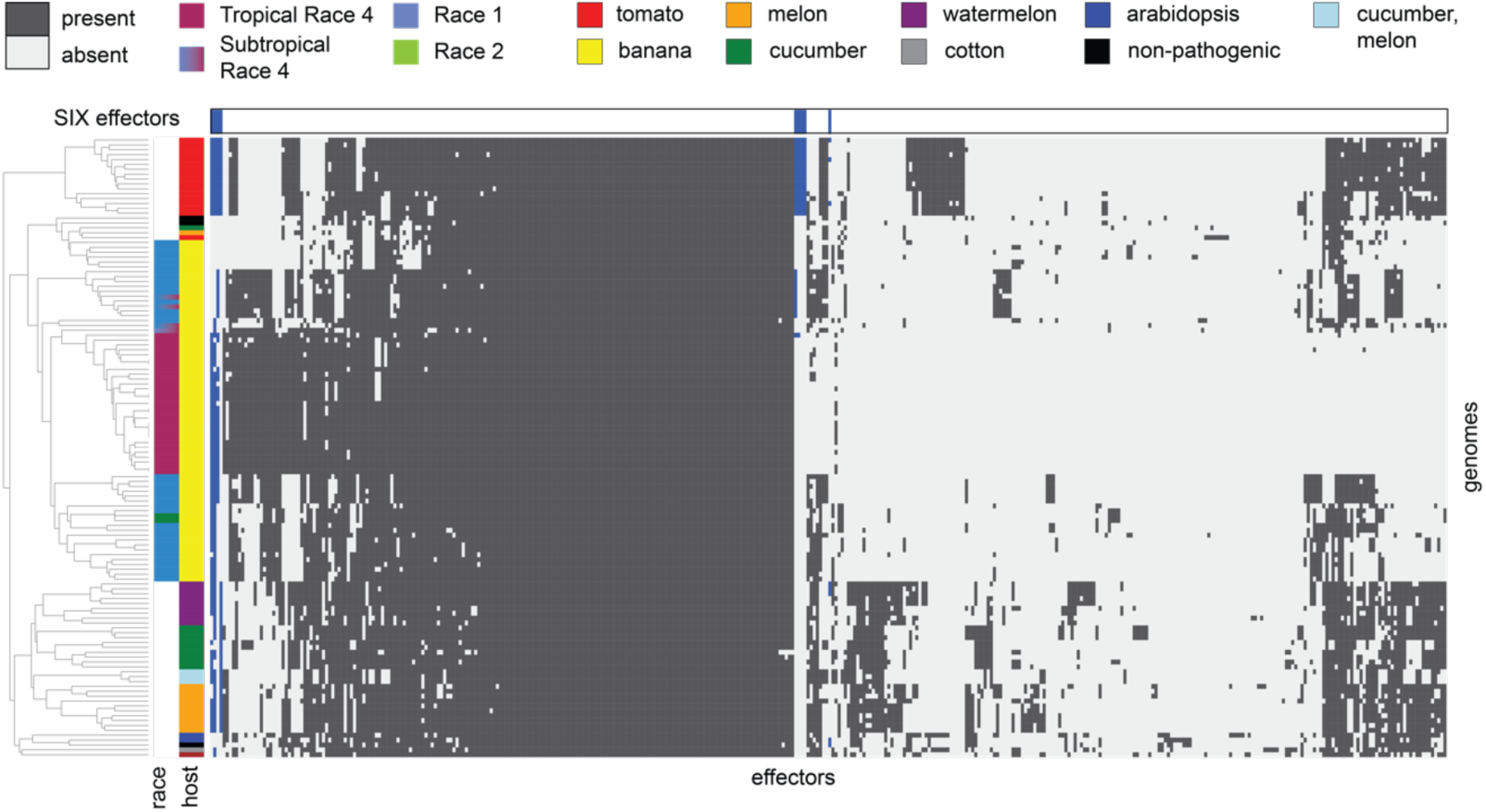
Predicted effector profile clusters *Fusarium* strains infecting the same host. *Fusarium* strains infecting banana (yellow) are separated from *Fusaria* infecting other hosts. In the set of 124 *Fusarium* isolates, 398 MIMP-associated effector families are predicted. The R1 strains (blue) have diverse effector profiles and do not cluster together, whereas the effector profiles of TR4 strains (purple) are highly similar. Nine SIX effectors (blue) are identified in the 398 MIMP-associated effector families and are present in Fusarium strains infecting tomato, from which they are originally identified, as well as in other Fusarium genomes.

### Recent segmental duplications drive accessory genome expansion in Fusarium oxysporum

We observed diverse ARs in *Fusarium* strains infecting banana with a high abundance of evolutionary young genes and genes that evolved via gene duplications (Fig. **4e**). Duplications have been previously reported in *F. oxysporum*, including *Fusarium* strains infecting banana (Kistler *et al*., 1995; Ma *et al*., 2010; Vlaardingerbroek *et al*., 2016; Armitage *et al*., 2018; Li *et al*., 2020). To better understand the role of gene duplications in the origin and diversification of ARs, we characterized gene duplications in *F. oxysporum* using MCscanX (Wang *et al*., 2012) and classified them into four categories (dispersed, proximal, tandem, and segmental) (Fig. **6a**). Remarkably, most genes in strain II5 (8,831 genes) have been affected by gene duplication during their evolution (Fig. **6b**, Supplementary Fig. 6). This high number of duplications include old gene duplications, recognized as distant homologs that acquired considerable number of mutations over time. When we apply a more stringent percent identity filtering, the number of duplicated genes is reduced, yet the number of segmental duplicated genes is least affected (supplementary table **4**), indicating that the segmental duplicated genes acquired less mutations and thus evolved more recently. We observed similar results for other *F. oxysporum* strains as well as for other *Fusarium* species; *F. verticillioides*, *F. graminearum,* and *F. solani* (syn. *Neocosmospora solani*), suggesting that duplications played a major role in *Fusarium* evolution (**Fig 6b**). Interestingly, we observed that segmental duplicated genes occur specifically in ARs (72 out of 124 in II5) and sub-telomeres (48 out of 124 in II5) in all chromosome-level *F. oxysporum* assemblies (Fig**. 6b**, Supplementary Fig. 6). In *F. solani*, which contains accessory chromosomes analogous to those observed in *F. oxysporum* (Coleman *et al*., 2009), 139 segmental duplicated genes were identified. In contrast, in *F. verticillioides* and *F. graminearum*, for which no accessory chromosomes have been described, no segmental duplicated genes are found (Fig. **6c-e**), underscoring the link between accessory chromosomes and segmental duplications in *Fusarium*. These duplications are possibly driven by TE activity (Wicker *et al*., 2010; Faino *et al*., 2016), and we observe that segmental duplications in TR4 strain II5 occur closer to transposable elements (median distance of 1734 kb, p <0.05, two sided Mann-Whitney U test) then other duplication types (dispersed 7721 bp, proximal 7503 bp, tandem 5941 bp), suggesting that TE activity influences the segmental duplications.

**Figure 6.**
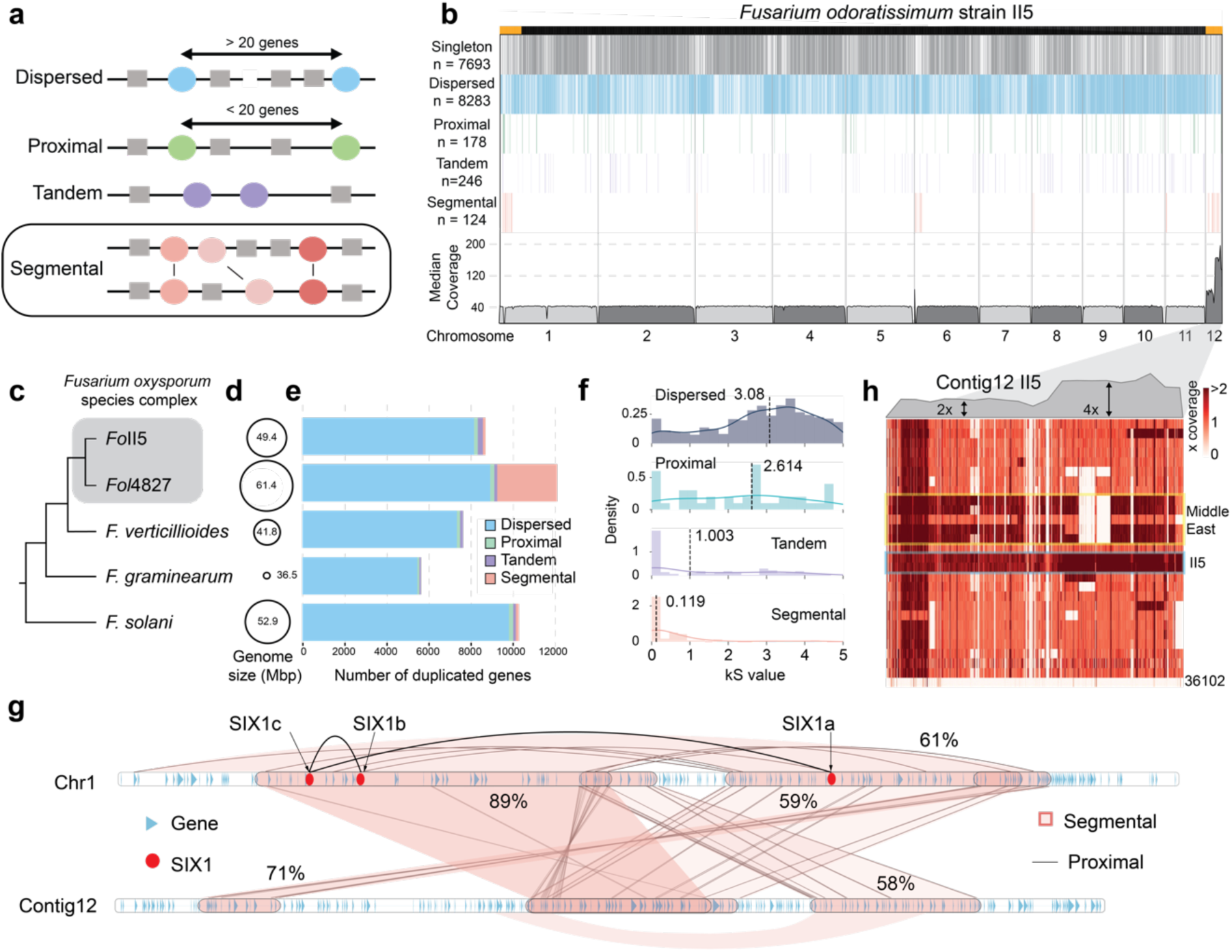
Segmental duplications are involved in the expansion of accessory regions in *F. oxysporum*. **a**) Four different types of duplications are distinguished. **b**) Segmental duplications (pink) occur mainly in the accessory regions in strain II5 (TR4). Sub-telomeric regions have more dispersed duplicated genes (blue) than the chromosomal arms. On average, the read coverage of II5 mapped onto the II5 reference genome is 40x, whereas contig 12 has a coverage between 80x and 160x, suggesting copy-number variations. **c**) The schematic tree depicts the relationship of *Fusarium* species with **d**) their genome sizes. **e**) Extensive segmental duplications are only found in *F. oxysporum* and *F. solani* and are virtually absent in *F. verticillioides* and *F. graminearum*. **f**) The age of duplications was estimated from the number of synonymous substitutions between duplicated gene pairs. Segmental duplications occurred most recently (kS = 0.119). kS distributions differ significantly between all duplication types (p <0.05, two sided Mann-Whitney U test). **g**) Genes encoding the virulence factor SIX1 (red) are part of a segmental duplication within chromosome 1. *SIX1a* and *SIX1b* are segmentally duplicated, and *SIX1c* is a proximal duplication sharing 82% amino acid sequence similarity to *SIX1a*. Light red blocks indicate segmental duplications, brown lines between the genes (blue arrows) highlight the genes involved in the segmental duplication. Not all genes in a block are part of the segmental duplication, for example a region on contig 12 is similar to the *SIX1* regions on chromosome 1, yet *SIX1* is absent on contig 12. **h)** Sequencing read coverage of strain II5 and three strains from the Middle East (JV11, ISR5, and JV14) show a two-fold increase relative to the genome-wide coverage. The other 26 strains do not show an increase in coverage, although smaller duplications are found throughout contig 12.

The highest number of segmental duplicated genes (2,887) is observed in *Fol*4287, carrying the largest accessory genome (19 Mb, 30% of the total genome size) (Fig. **6d**) and 2,366 (82%) of the segmental duplicated genes are in ARs. Interestingly, many segmental duplicated genes are shared between accessory chromosomes 3 and 6 (Supplementary Fig. 7), as previously observed (Ma *et al*., 2010), indicating that accessory chromosomes evolved by inter-chromosomal segmental duplication that drive the expansion of ARs in *Fusarium*.

Duplications of AR genes significantly effect effector repertoires as 336 out of 669 predicted effectors in II5 evolved via duplications, and nine effectors are part of segmental duplications. Among the segmental duplicated effectors is *SIX1*, an effector that is essential for full virulence to banana (Widinugraheni *et al*., 2018). Three copies of *SIX1* (*SIX1a*,*b*,*c*) are present in the AR on chromosome 1 of the TR4 strains M1, 36102 and II5. *SIX1a* and *SIX1b* are part of a segmentally duplicated block (Fig. **6g**), whereas *SIX1c* is a proximal duplication that shares 82% amino acid sequence identity with *SIX1a* and 73% sequence identity with *SIX1b*. In contrast, the R1 strain CR1.1 contains only one copy of *SIX1*, resembling *SIX1c*, on accessory contig 12. The duplicated blocks with *SIX1* are interspersed with non-duplicated genes (Fig. **6g**), indicating that over time these blocks further diverged by gene gains and losses. Interestingly, the segmentally duplicated *SIX1* blocks also share similarity to a region on contig 12 (Fig. **6g**), yet no copy of *SIX1* is present, and thus *SIX1* is either lost or has been gained on chromosome 1 after the segmental duplication between chromosome 1 and contig 12 occurred. Segmental duplications therefore contribute to the evolution of virulence factors such as *SIX1*, which is essential for successful banana infection (Widinugraheni *et al*., 2018).

To estimate the relative timing of the duplication events, we used the synonymous substitution rate (dS) between duplicated gene pairs as a proxy of time. Dispersed duplicated genes had the highest average dS value (3.08), suggesting that these are the most ancient duplicates, while genes pairs that arose via segmental duplication evolved more recently, with a significantly lower average dS value of 0.61 (Fig. **6f**). To assess if even more recent large-scale duplications are present in our global panel, we determined the read depth of all isolates sequenced with short reads against II5. Based on the genome-wide average read depth (∼40x coverage), we determined that contig 12 has been entirely duplicated (read depth of ∼80x coverage), while a section (position ∼64-116 kb; ∼160x coverage) occurs four times (Fig. **6h**). This demonstrates that the II5 carries two (nearly) identical copies of contig 12 (i.e., aneuploidy), in addition to the previously observed segmental duplications (Fig. **6h**). No copy number variation of the accessory region of chromosome 1 is observed. The four partial copies of contig 12 possibly originate from an additional duplication, it remains unclear if this duplicated stretch is attached to a chromosome, or if it possibly gave rise to an additional smaller accessory chromosome. To resolve the origin of this duplicated region, we mapped the long reads to the assembly. We observe many clipped reads in this region, which do not entirely correspond to the contig 12 assembly, however the clipped regions do not align to another chromosome. This suggests the region is not attached to another chromosomal arm, leaving the possibility that the duplicated region could represent a duplicated chromosomal arm. We didn’t observe a threefold increase on contig 12, suggesting that the duplication happened multiple times. Interestingly, aneuploidy of contig 12 can also be observed in three additional TR4 strains from the Middle East, but not in the other 26 TR4 strains (Fig. **6h**). Collectively, we demonstrate that the evolutionary dynamics of ARs in *F. oxysporum* is driven by recent segmental duplications and aneuploidy.

## Discussion

Many fungal plant pathogens evolved compartmentalized genomes with conserved core regions and variable accessory regions (ARs) (Dong *et al*., 2015; Torres *et al*., 2020). Some members of the *Fusarium oxysporum* species complex are important plant pathogens and are well-known to have extensive ARs that can encompass entire chromosomes, which have been linked to pathogenicity towards specific hosts (Sánchez-Vallet *et al*., 2018). Here we use a pan-genomic framework to study the occurrence, composition, and evolution of ARs in a global collection of banana infecting *Fusarium* species. Strains carry up to 15 Mb of ARs that contain *in planta* induced effector genes as well as most *SIX* effectors. ARs are highly divergent among strains and races, and consequently, we cannot identify shared ARs or accessory chromosomes that can be linked to pathogenicity towards banana. We furthermore demonstrate that *Fusarium* evolved by extensive duplications and that ARs specifically are shaped by recent segmental duplications in *F. oxysporum* and *F. solani*. Lastly, we uncover that an accessory contig in II5 underwent recent aneuploidy, suggesting that these processes drive the emergence and evolution of ARs in *F. oxysporum*.

Fusarium wilt of banana is a major threat to food security as banana is a major staple crop in tropical and subtropical regions (Drenth & Kema, 2021; van Westerhoven *et al*., 2022b). Compared to R1 strains, which caused epidemics in Gros Michel bananas in the 1900s, TR4 strains emerged more recently (1967) and cause the ongoing FWB pandemic in Cavendish plantations (van Westerhoven *et al*., 2022b). TR4 strains are genetically highly similar, corroborating that a single clone that circumvents Cavendish resistance drives the ongoing pandemic (Ordonez *et al*., 2015; van Westerhoven *et al*., 2022b). By contrast, R1 strains are genetically diverse, which is possibly due to prolonged co-evolution of R1 strains with a plethora of genetically diverse banana varieties in the center of origin in Southeast Asia (Perrier *et al*., 2011). Extensive genetic diversity among *Fusarium* strains that infect the same host is not uncommon (Baayen *et al*., 2000; van Dam *et al*., 2016; Henry *et al*., 2021; Fayyaz *et al*., 2023), yet their ARs and effector profiles are typically remarkably similar (van Dam *et al*., 2016; Fokkens *et al*., 2018; Brenes Guallar *et al*., 2022). We, however, observe that the ARs and effector profile of R1 strains show extensive variation, underscoring their diversity and possibly suggesting that different mechanisms contribute to disease in banana (Henry *et al*., 2021; Brenes Guallar *et al*., 2022). *Fusarium* strains that infect strawberry have two distinct ARs, one that contributes to yellowing and the other to wilting (Henry *et al*., 2021). Quantitative differences in banana corm-discoloration have been noted between R1 strains (Garcia-Bastidas, 2019), and our results suggest that this might relate to differences in effector profiles, yet further functional assays are needed to elucidate the link between specific effectors and phenotypic variation.

We identify abundant and recent segmental duplications in *F. oxysporum* ARs, as well as aneuploidy of contig 12 in TR4 strain II5. Previous studies have observed large-scale duplications and deletions in the ARs of *F. oxysporum* during *in vitro* chromosome transfer (Vlaardingerbroek *et al*., 2016; Li *et al*., 2020), yet our results show that duplication of ARs also occur in *Fusarium* strains sampled directly from infected banana plants. Although variations in chromosome number typically comes with a fitness cost (Todd *et al*., 2017), it can also provide genetic variation necessary for adaptation, for example increasing virulence or fungicide resistance (Sionov *et al*., 2010; Ropars *et al*., 2018). We speculate that the observed large-scale duplications play a major role in the origin and evolution of ARs in *Fusarium*. In the wheat pathogen *Zymoseptoria tritici*, chromosomal duplication, together with breakage and fusion, shape the evolutionary dynamics of accessory regions (Croll *et al*., 2013; Möller *et al*., 2018). Similar dynamics of chromosome duplication, breakage, and fusion may underly the recent segmental duplications observed in *Fusarium*. For example, the fusion of a (partially) duplicated contig 12 with the arm of chromosome 1 in strain II5 could explain the observed homology between the ARs. The process underlying these chromosome dynamics remains unclear. *F. oxysporum* has long been considered to evolve strictly asexually (Gordon & Martyn, 1997), however, the presence of an active sexual cycle in *F. oxysporum* populations has recently been proposed (McTaggart *et al*., 2022; Fayyaz *et al*., 2023) and we similarly observe reticulation between strains that caused FWB (Supplementary Fig. 8), which supports an evolutionary history in which ancient or infrequent sexual recombination is followed by clonal expansion of selected lineages. In this scenario, incomplete chromosome pairing and non-disjunction during meiosis might drive aneuploidy and ARs diversification (Fayyaz *et al*., 2023). We show that ARs are similar within individual *Fusarium* lineages but are genetically distinct when comparing strains from genetically distant lineages. Individual lineages accumulate genomic changes over time, and this process might be sufficient to explain ARs diversity. If new *Fusarium* lineages indeed arise from a meiotic cycle followed by clonal expansion, recombination, gain, or loss of ARs during meiosis might further explain the diversity of ARs between lineages. Similarly, the absence of meiotic recombination during subsequent clonal expansion would explain the similarity of ARs within clonal lineages. However, we observe aneuploidy within individual clonal lineages, for instance the duplication of contig 12 in some *Fusarium* strains, which demonstrates that aneuploidy can occur independent of a meiotic cycle. Thus, chromosome dynamics in *F. oxysporum* require further elucidation and, so far, the mechanisms leading to chromosomal duplications are unclear; they might arise from non-disjunction during mitosis or meiosis (Fayyaz *et al*., 2023), from incomplete chromosome loss following heterokaryon formation (Harrison *et al*., 2014; Shahi *et al*., 2016), or through consecutive horizontal chromosome transfers when strains acquire the identical chromosome several times.

In contrast to the structural variation in the ARs, the core chromosomes in *Fusarium* are remarkably stable. Although ARs are likely more tolerant to structural variation as they encode fewer essential genes, additional processes can influence genome stability. First of all, the accessory genome is enriched in transposable elements (Figure 2b) and the activity of these elements can contribute to structural variation including gene duplications (Faino *et al*., 2016; Torres *et al*., 2021; Wang *et al*., 2022; Stalder *et al*., 2022). Possibly the activity of transposable elements in the accessory genomic regions gave rise to the observed segmental duplications, for example, in plant genomes large-scale duplications can arise through double stranded break repair upon TE-induced double stranded breaks (Wicker *et al*., 2010; Wang *et al*., 2022). For example, the presence of histone modifications such as trimethylation of lysine 27 on histone 3 (H3K27me3), a histone modification often enriched in ARs (Fokkens *et al*., 2018; Möller *et al*., 2019; Torres *et al*., 2020) in fungal plant pathogens, has been linked to genome instability (Seidl *et al*., 2016; Möller *et al*., 2019). H3K27me3 also occurs in the ARs of *F. oxysporum* infecting tomato (*Fol*4287), but can also be found in smaller core chromosomes (‘fast-core’ chromosomes, 9 – 11 (Fokkens *et al*., 2018)) that do not seem to undergo segmental duplications or aneuploidy, and thus additional factors likely influence chromosome stability in *F. oxysporum*.

Extensive duplications similarly occur in other plant pathogens (Dutheil *et al*., 2016; Müller *et al*., 2019; Wyka *et al*., 2021; Wacker *et al*., 2023), and are thought to be important drivers in co-evolutionary arms races with their hosts. Understanding the evolution of ARs in *Fusarium* can facilitate the discovery of new effector genes and provide insights into effector diversification. Knowledge of effector profiles is crucial for designing effective disease control strategies and supports the identification of durable resistances in crops (Vleeshouwers *et al*., 2008).

## Acknowledgments

ACW, GHJK, CAG, GNT, RS and JD were supported by the Bill and Melinda Gates Foundation, grant number AG - 4425. The funders had no role in study design, data collection and analysis, decision to publish, or preparation of the manuscript. We thank the Research Support Group (RSG) at KeyGene (Wageningen, The Netherlands), which provided DNA isolation, library preparation and long-read sequencing for three of the strains. Banana research at Wageningen University has been supported by the Dutch Dioraphte Foundation, grant number 20 04 04 02.

## Notes

### Competing Interest Statement

The authors have declared no competing interest.

### Summary of Updates

Updated manuscript based on reviewers' feedback; Figure 1-6 revised, S-files updated.

