## Supplementary Information for "Segmental Duplications Drive the Evolution of Accessory Regions in a Major Crop Pathogen"

***New Phytologist* Supporting Information**

Article acceptance date: Click here to enter a date.

The following Supporting Information is available for this article:

**This file includes:**

Supporting material and methods
Figures S1 to S8
Tables S2 to S4

**Other supporting materials for this manuscript include the following:**

Table S1
Table S5

**Methods S1**

**DNA extraction and sequencing**

**Protocol Nanopore sequencing for isolate 36102, II5, and CR1.1**

For three banana infecting *Fusarium* strains (II5, 36102, and CR1.1), Oxford Nanopore sequencing (Oxford Nanopore Technologies, Oxford, UK) was performed by KeyGene (the Netherlands). DNA was isolates as described by Garcia-Bastidas et al.[1].The isolates were cultured on potato dextrose agar (PDA; Sigma-Aldrich, Zwijndrecht, the Netherlands) plates at 28°C. For DNA extraction, 0.5 cm2 agar blocks colonized with *Fusarium* were used to inoculate 400 mL liquid Mung bean media. The cultures were incubated at 25°C and 130 rpm for four days. Conidia were collected by filtering and centrifugation and washed twice with demineralized water followed by two washes with 500 mM $NaCl_2$/ 50 mM EDTA pH 8.0. After washing, the conidia were collected, and the excess of moisture was squeezed out. The dried tissue was collected and stored at -80°C for later use.

Deep-frozen conidia were ground to powder in liquid nitrogen. The powder was resuspended in four volumes of DNA extraction buffer (100 mM tris-HCl pH8, 70 mM EDTA pH8.0, 2% (v/v) SLS, 2% (v/v) 2-mercaptoethanol, 100 µg/mL proteinase K) and the homogenate was incubated at 55°C for 1 h. After incubation, the homogenate was centrifuged for 10 min at 5,087 g and the supernatant was extracted with 1 volume of chloroform / isoamyl alcohol (24:1). Finally, DNA was precipitated with isopropanol, air-dried, and resuspended in nuclease-free water. DNA quantity and purity was determined with the Nanodrop UV-Vis spectrophotometer (Thermo Fisher Scientific, Wilmington, NC, USA) and the Qubit 2.0 fluorometer (Thermo Fisher Scientific, Waltham, MA, USA). Integrity assessment of the genomic DNA was performed using a Femto Pulse system (Agilent Technologies, Santa Clara, CA, USA).

Short DNA fragments (≤25 kb) were removed using Blue Pippin (Sage Sciences, Beverly, MA, USA). For each strain, a DNA sequencing library was prepared with the 1D ligation Sequencing Kit (SQK-LSK109) and subsequently sequenced with one PromethION (R9.4.1) flow cell and base calling was performed on the instrument using MinKNOW version 3.4.6.

**Protocol Nanopore sequencing, isolates from Cuba and Mozambique**

High-molecular weight DNA isolation was made according to protocol described by Chavarro-Carrero et al.[2], with some modifications. Conidia were harvested from PDA and subsequently grown for three days in potato dextrose broth (PDB, 1/4 strength) at 25°C and 150 rpm. Fungal material was collected on Miracloth, freeze-dried overnight and ground to powder. DNA was extracted by incubating 300 mg of fungal material for 1 h at 65°C with 350 µl DNA extraction buffer (0.35 M Sorbitol, 0.1 M Tris-base, 5 mM EDTA pH 7.5), 350 µl nucleic lysis buffer (0.2 M Tris, 0.05 M EDTA, 2 M NaCl, 2% CTAB) and 162.5 µl Sarkosyl (10% w/v) with 1% β-mercaptoethanol. Next, 400 µl of phenol/chloroform/isoamyl alcohol (25:24:1) was added, shaken, and incubated at room temperature (RT) for 5 min before centrifugation at 16 000 g for 15 min. The aqueous phase was transferred to a tube and 10 µl of RNase (10 mg $\mu L^{-1}$) was added and incubated at 37°C for 1 h. Supernatant was extracted using half a volume of chloroform and centrifuged at 16,000 g for 5 min at RT, after which the chloroform extraction was repeated. Next, the aqueous phase was mixed with 10 volumes of 100% ice-cold ethanol, incubated for 30 min at RT, and the DNA was fished out, transferred, and washed twice with 500 µl 70% ethanol. Finally, the DNA was air-dried, resuspended in nuclease-free water, and incubated at 4°C for 2 days. DNA quantity and purity was determined with the Nanodrop UV-Vis spectrophotometer (Thermo Fisher Scientific, Wilmington, NC, USA) and the Qubit 2.0 fluorometer (Thermo Fisher Scientific, Waltham, MA, USA).

Library preparation with the Rapid Sequencing Kit (SQK-RAD004) was performed according to the manufacturer's instructions (Oxford Nanopore Technologies, Oxford, UK) with 400 ng HMW DNA. An R9.4.1 flow cell (Oxford Nanopore Technologies, Oxford, UK) was loaded and run for 24 h. Base calling was performed using Guppy (version 3.1.5; Oxford Nanopore Technologies, Oxford, UK) with the high-accuracy base-calling algorithm.

**Protocol short-read sequencing**

Monosporic cultures were grown in PDB and incubated under continuous shaking (125 rpm) at RT. After seven days of incubation, fungal biomass was collected by filtering the cultures through cheesecloth and samples were freeze-dried in a 2 mL tube for 21 h. The genomic DNA of each isolate was extracted using the DNA-Kit Wizard Magnetic DNA Purification System for Food kit (Promega, USA) or the The MasterPure™ Yeast DNA Purification (LGC Biosearch Technologies), according with the manufacturer protocol. DNA concentration was quantified using the Quant-iT™ PicoGreen™ dsDNA Reagent Invitrogen in a TECAN 2000 Analyzer, (Thermo Fisher Scientific, Switzerland), and the quality was checked on an agarose gel. DNA samples of high quantity and quality (500 ng; 100 ng/uL) of each isolate were used for sequencing.

**Supplementary Table 2.** Statistics of the long-read nanopore sequencing data of seven Fusarium strains.

| | **CR1.1 (R1)** | **C058 (R1)** | **C135 (R2)** | **C081 (R2)** | **II5 (TR4)** | **36102 (TR4)** | **M1 (TR4)** |
|---|---|---|---|---|---|---|---|
| **Number of reads** | 2.089,010 | 1,156,301 | 690,046 | 836,000 | 34,604,616 | 2,062,327 | 261,898 |
| **Mean read length (bp)** | 21,918 | 4,020 | 6,826 | 5,602 | 28,940 | 34,932 | 6,858 |
| **Median read Length (bp)** | 14,352 | 2,300 | 4,067 | 3,421 | 28,941 | 35,828 | 3,087 |
| **Read N50 (bp)** | 39,394 | 7,364 | 12,831 | 10,355 | 37,836 | 46,492 | 16,112 |
| **Mean read quality** | 10.50 | 9.10 | 9.30 | 9.40 | 10.40 | 10.50 | 7.70 |
| **Median read quality** | 10.70 | 11.70 | 11.40 | 11.60 | 10.60 | 10.70 | 8.30 |
| **Bases (Mb)** | 45,787 | 4,648 | 4,710 | 4,683 | 100,149 | 72,042 | 1,796 |

**Supplementary Table 3.** Genome assembly statistics of seven Fusarium strains sequenced with nanopore long-read sequencing technology.

| | **CR1.1 (R1)** | **C058 (R1)** | **C135 (R2)** | **C081 (R2)** | **II5 (TR4)** | **36102 (TR4)** | **M1 (TR4)** |
|---|---|---|---|---|---|---|---|
| **Contigs** | 12 | 13 | 15 | 15 | 12 | 11 | 12 |
| **N50 (bp)** | 4,611,249 | 4,548,202 | 4,717,159 | 4,662,367 | 4,544,333 | 4,861,540 | 4,516,955 |
| **Telomeres** | 19 (of 24) | 24 (of 26) | 21 (of 30) | 25 (of 30) | 20 (of 24) | 21 (of 22) | 20 (of 24) |
| **Length (Mb)** | 48.0 | 50.1 | 52.3 | 50.8 | 48.6 | 45.8 | 47.7 |

**Supplementary Table 4.** Number of duplicated genes as reported compared to the number of duplicated genes using a stricter filtering approach.

| | Gene duplications (reported) | Gene duplications (>50% identity) | Percentage genes removed with strict filtering |
|---|---|---|---|
| Filtering settings | 50% query coverage, >20% identity | 50% query coverage, >50% identity | |
| Dispersed | 8,283 | 2,354 | 72% |
| Proximal | 178 | 26 | 85% |
| Tandem | 246 | 108 | 56% |
| Segmental | 124 | 74 | 40% |
| Total duplicates | 8,831 | 2,562 | 70% |

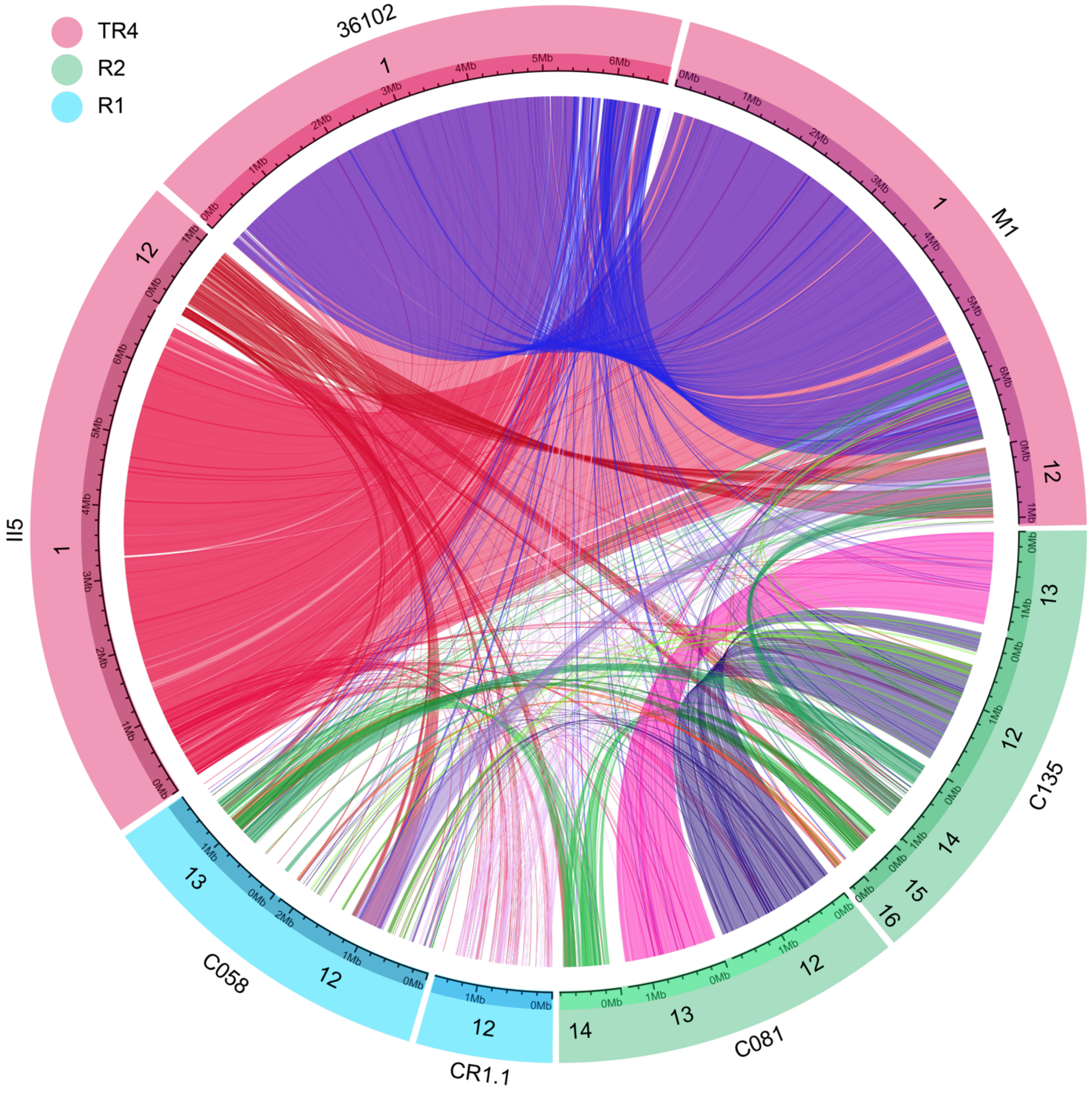


**Supplementary Figure 1. Nucleotide alignments between ARs of the chromosome level genome assemblies identify diverse ARs.** ARs from the seven chromosome-level *Fusarium* strains infecting banana are identified based on a lack of co-linearity in with *Fusarium oxysporum* f. sp. *lycopersici Fol*4287. Ribbons depict nucleotide alignments between two regions. *Fusarium* strain II5 and M1 (both TR4) share the ARs on chromosome 1 and on contig 12, while strain 36102 (TR4) carries only the AR on chromosome 1. R2 strains C135 and C081 share two accessory chromosomes with little similarity to any other ARs. Some links are present between all ARs, indicating smaller shared genomic regions, however Race 1 strains do not share any larger AR.

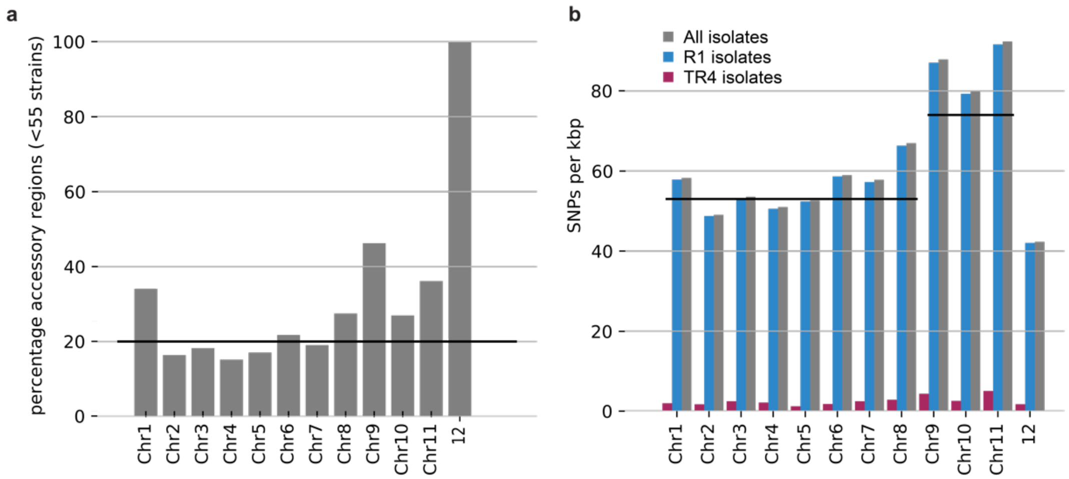


**Supplementary Figure 2 - Chromosome 9, 10, and 11 of *Fusarium* strain II5 (TR4) share few genomic regions with other *Fusarium* strains infecting banana and have more polymorphic sites.** **a**) Contig 12 occurs in less than 55 of the *Fusarium* strains infecting banana and is entirely composed of accessory regions (ARs). Chromosomes 9, 10 and 11 have more ARs than the genome-wide average (20.2%, black line). **b**) Single nucleotide polymorphisms (SNPs) in the genome of II5 (TR4). TR4 strains are less diverse and show on average 1.9 SNPs per Kbp (red bars), while R1 strains are more distantly related and encode on average 57.5 SNPs per Kbp (blue bars). Chromosomes 9, 10, and 11 contain more SNPs compared with the other chromosomes (74 SNPs per Kbp vs 53 SNPs per Kbp, indicated by the horizontal black lines).

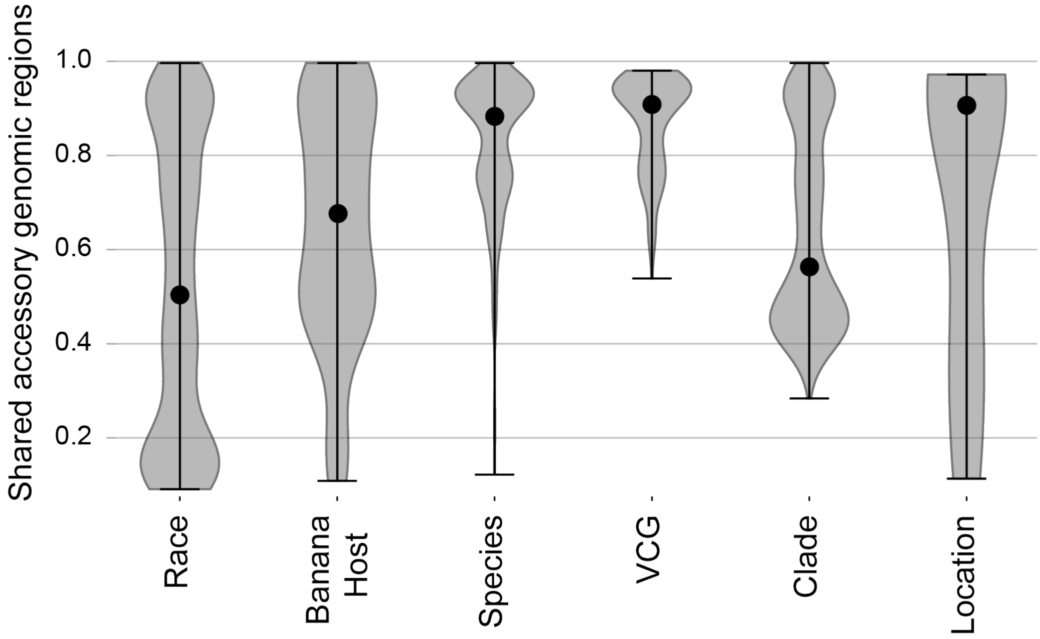


**Supplementary Figure 1. Fraction of shared accessory regions (ARs) in different subgroups of *Fusarium* strains infecting banana.** Strains are grouped per race (Race), banana host from which the strain has been isolated (Banana Host), phylogenetic species (Species), vegetative compatibility group (VCG), phylogenetic clade (Clade), or origin of isolation (Location). Species and Clade are obtained from the phylogenetic tree, whereas VCG has been previously determined based on compatibility assays. Per group, the pairwise fraction of shared ARs is calculated. The violin plots show the distribution of each pairwise comparison, with low values indicating that some strains belonging to the same category share little AGRs and black dots highlight the median fraction. Strains belonging to the same species (median = 0.88, black dot) and VCG (median

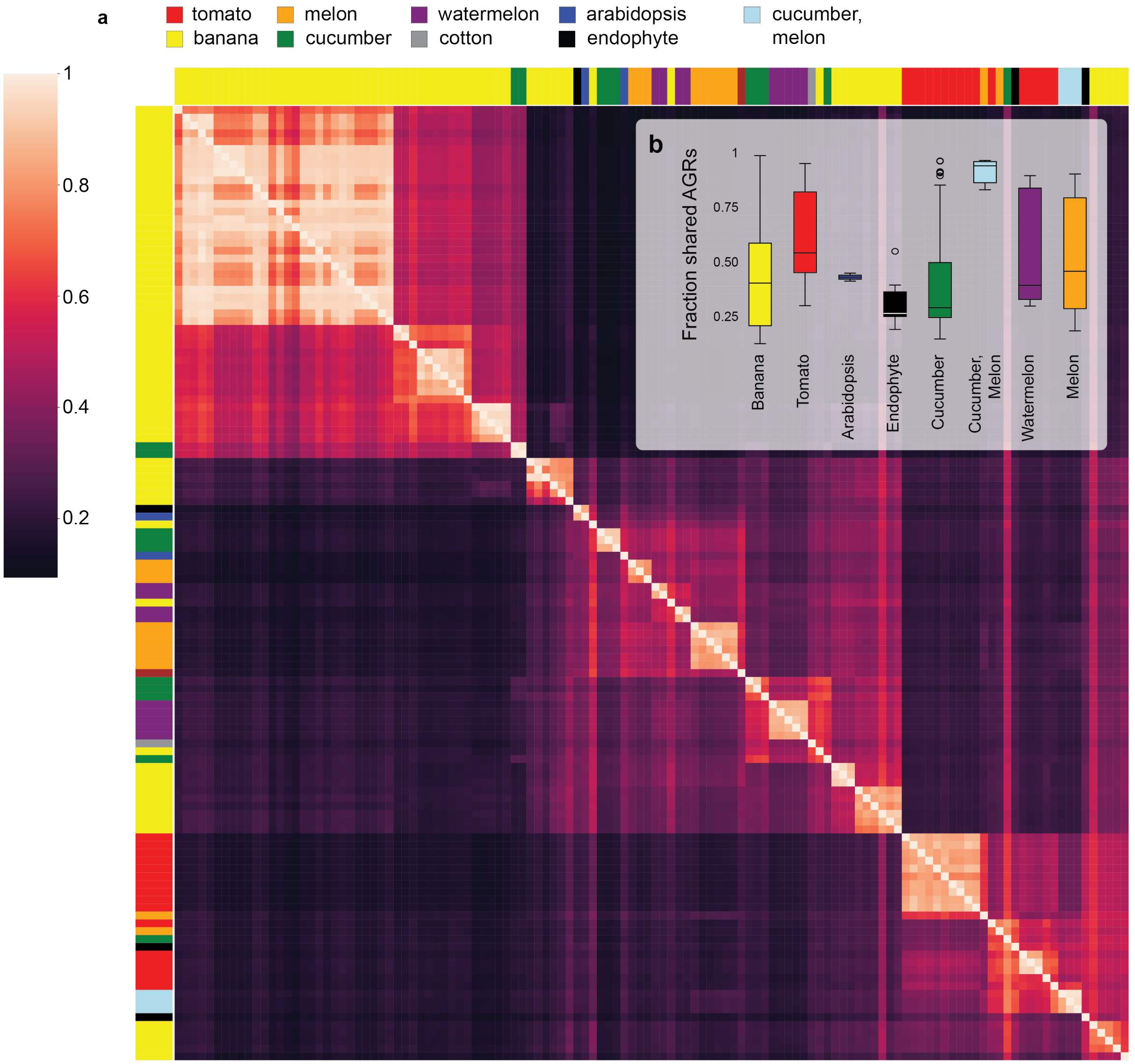


**Supplementary Figure 2. Accessory regions (ARs) are shared between genetically related *Fusarium* strains. a)** Heatmap shows the fraction of shared ARs between *Fusarium* strains infecting different host plants, ordered by their phylogenetic relatedness. The fraction is calculated relative to the amount of ARs in the strain on the x-axis, resulting in light lines (high similarity) in strains with little ARs. **b**) Strains infecting the same host can share little to no ARs, for example, as observed in banana or cucumber, while other strains such as those infecting tomato always share at least 30% of their ARs.

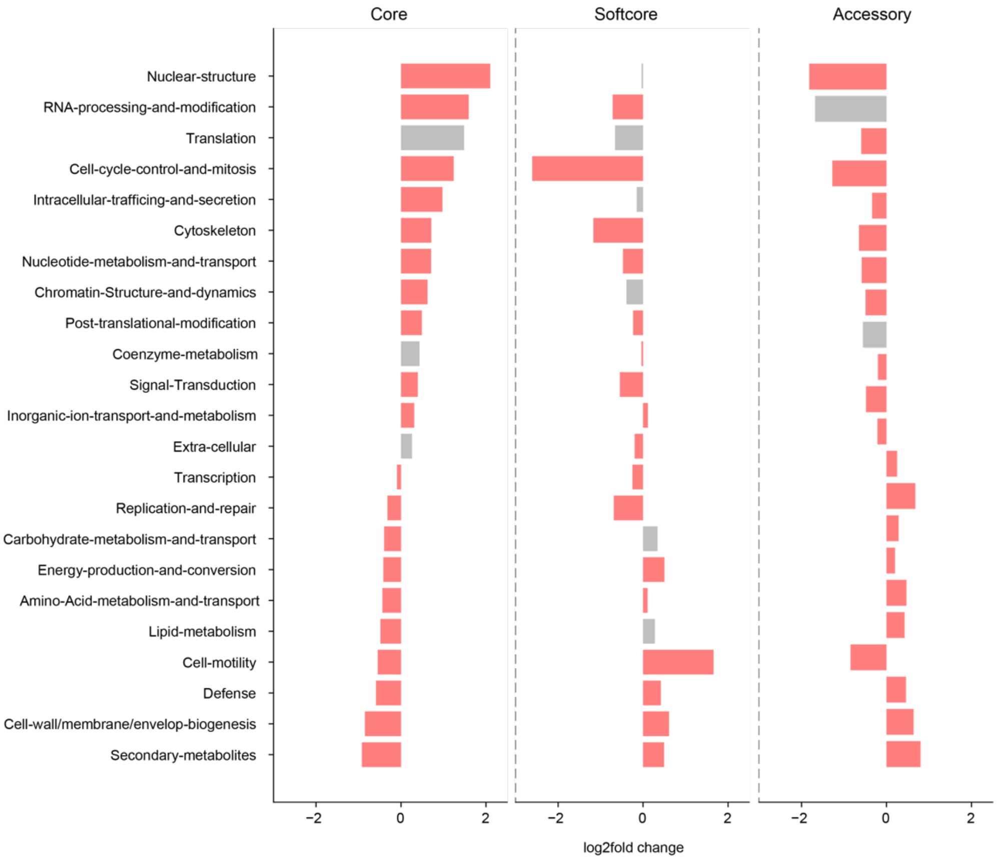


**Supplementary Figure 3. Functional enrichment of core, softcore and accessory genes.** The bars show the $\log_2$-fold change per gene function in the core, softcore, and accessory genes. Red color indicates a significant $\log_2$ fold-change for the gene category (P-value < 0.05, Fisher's exact test). Conserved core genes are enriched for housekeeping functions, whilst accessory genes are enriched in defense compounds and secondary metabolites.

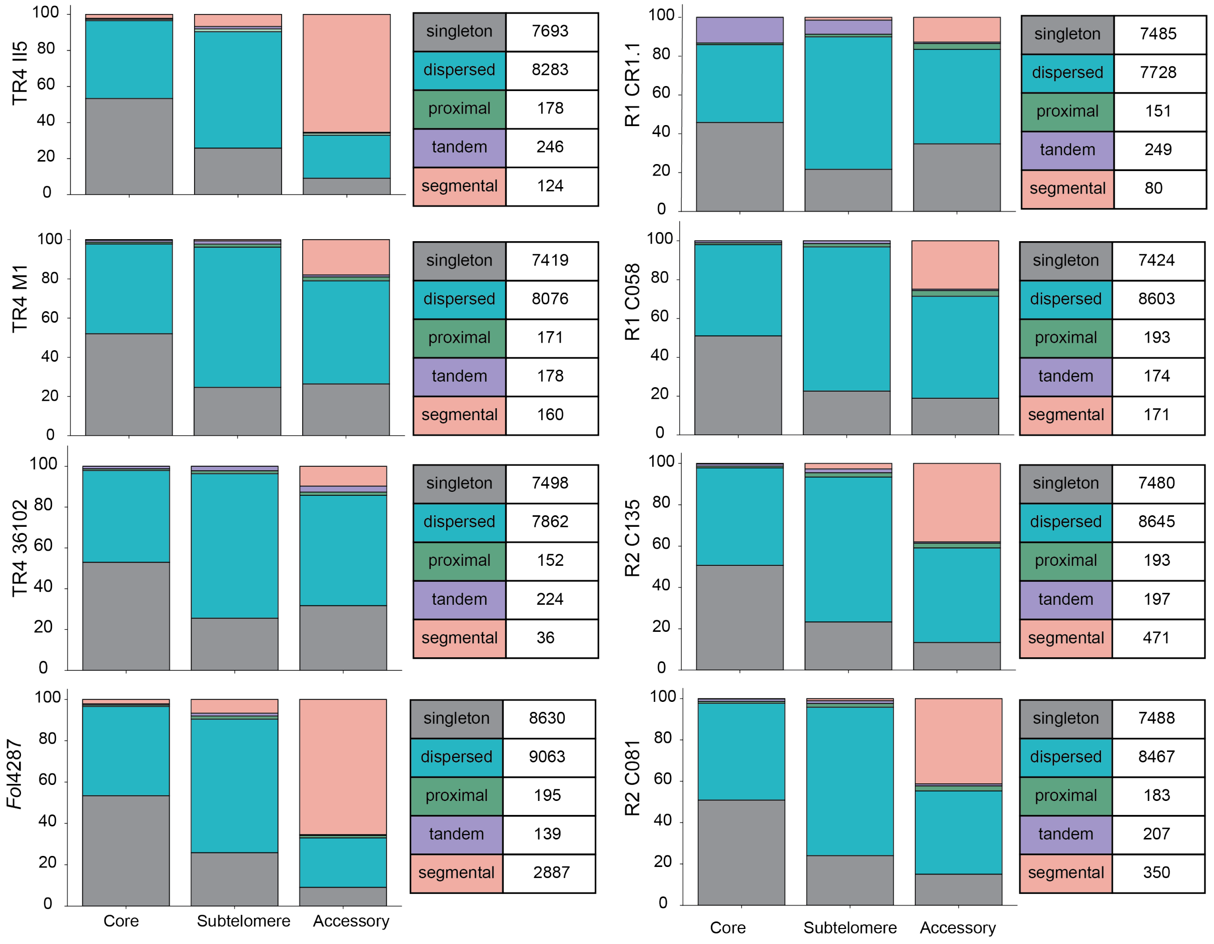


**Supplementary Figure 4. Duplication types found in the core, softcore and accessory regions in the seven chromosome level genome assemblies of Fusarium infecting banana and the *Fusarium* strain infecting tomato (*Fol*4287).** Segmental duplications are most abundant in accessory regions (ARs). The percentage and number of gene duplications identified and classified by MCScanX are shown for the seven chromosome-level genome assemblies of Fusarium strains infecting banana as well as of *Fusarium oxysporum* f.sp. *lycopersici* (*Fol*4287). Singletons are most abundant in the core genome, yet gene duplications occur throughout the genome, especially in sub-telomeric and ARs. While core regions have little segmental duplications, the ARs show extensive segmental duplication. The number of segmental duplications varies between Fusarium strains, and most segmental duplications can be found in *Fol*4287.

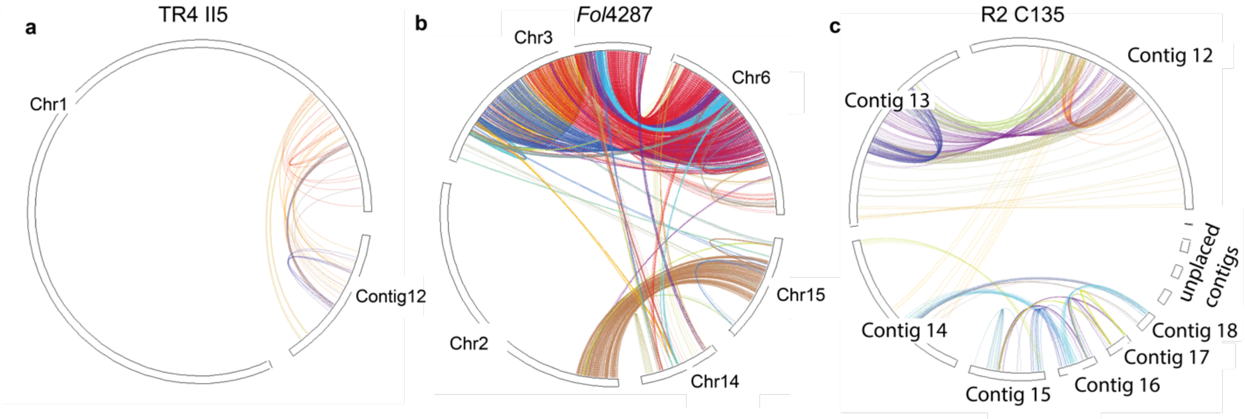


**Supplementary Figure 5.** Segmental duplications between accessory regions. Ribbons highlight gene pairs that are part of segmental duplications. In *Fol*4287 (b) chromosome 3 and chromosome 6 share large segmental duplications. Similarly, *Fusarium* strain C135 (c) shows many segmental duplications between contig 12 and contig 13.

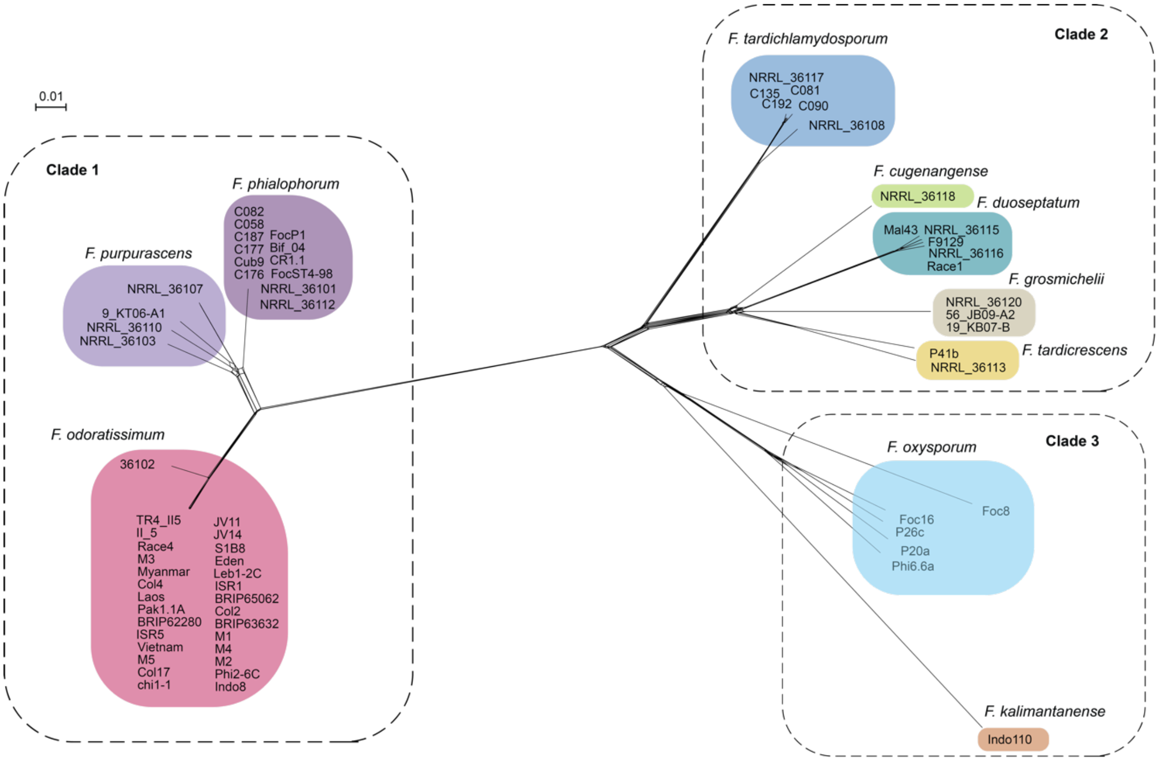


**Supplementary Figure 8.** Splitstree network shows indications of reticulation that might suggest recombination between some *Fusarium* strains infecting banana. The network is based on single nucleotide polymorphism found in 69 strains compared with the reference strain II5, excluding the accessory regions on Chromosome 1 and Contig 12. Most recombination signal is present in the branches leading to clade 2 and to *F. purpurascens* and *F. phialophorum*.
